# Genetic basis of body shape variation along the benthic-pelagic axis in cichlid fishes

**DOI:** 10.1101/2021.10.02.462884

**Authors:** Leah DeLorenzo, Destiny Mathews, A. Allyson Brandon, Mansi Joglekar, Aldo Carmona Baez, Emily C. Moore, Patrick J. Ciccotto, Natalie B. Roberts, Reade B. Roberts, Kara E. Powder

## Abstract

Divergence along the benthic-pelagic axis is one of the most widespread and repeated patterns of morphological variation in fishes, producing body shape diversity associated with ecology and swimming mechanics. This ecological shift is also the first stage of the explosive adaptive radiation of cichlid fishes in the East African Rift Lakes. We use two hybrid crosses of cichlids (*Metriaclima* sp. x *Aulonocara* sp. and *Labidochromis* sp. x *Labeotropheus* sp., >975 animals total) along the benthic-pelagic ecomorphological axis to determine the genetic basis of body shape diversification. Using a series of both linear and geometric shape measurements, we identify 55 quantitative trait loci (QTL) that underlie various aspects of body shape variation associated with benthic-pelagic divergence. These QTL are spread throughout the genome, each explain 3.0-7.2% of phenotypic variation, and are largely modular. Further, QTL are distinct both between these two crosses of Lake Malawi cichlids and compared to previously identified QTL for body shape in fishes such as sticklebacks. We find that body shape is controlled by many genes of small effects. In all, we find that convergent benthic and pelagic body phenotypes commonly observed across fish clades are most likely due to distinct genetic and molecular mechanisms.

## INTRODUCTION

Body shape variation is common across all vertebrates and has important consequences for an animal’s ecology, locomotion, thermodynamics, and even speciation (Arnold, 1983, 1992; Coyne & Orr, 2004; S.T. Friedman, Price, & Wainwright, 2021; Ruff, 1991; Schluter, 1996a, 2000; Smith, Nelson-Maney, Parsons, Cooper, & Albertson, 2015). A major axis of this morphological variation is body elongation, which occurs within reptiles (Bergmann & Irschick, 2012; Losos, 2009; Wiens & Slingluff, 2001), carnivorous mammals (Law, 2019, 2021), and fishes (S. T. Friedman et al., 2020; Price et al., 2019; Ward & Mehta, 2010). Within fishes, elongation of the body is associated with a major ecomorphological divergence, the benthic-pelagic axis.

The benthic-pelagic axis is one of the most widespread and repeated patterns of morphological variation within both marine and freshwater fishes, affecting traits such as body shape, fin position and musculature, and jaw mechanics (Burns & Sidlauskas, 2018; Cooper, Carter, Conith, Rice, & Westneat, 2017; Cooper et al., 2010; Hatfield & Schluter, 1999; Hulsey, Roberts, Loh, Rupp, & Streelman, 2013; Kusche, Recknagel, Elmer, & Meyer, 2014; Muschick, Indermaur, & Salzburger, 2012; Price et al., 2019; Ribeiro, Davis, Rivero-Vega, Orti, & Betancur, 2018; Robinson & Wilson, 1994; Rogers & Jamniczky, 2014; Schluter, 1996b; Walker, 1997; Willacker, Von Hippel, Wilton, & Walton, 2010). This ecological and morphological divergence occurs similarly in ancient fish radiations (Ribeiro et al., 2018), at the macroevolutionary level (Claverie & Wainwright, 2014; S. T. Friedman et al., 2020; Larouche et al., 2020; Price et al., 2019), and repeatedly at the microevolutionary level. Benthic-pelagic body shape variation occurs during both sympatric and allopatric speciation, for instance in sticklebacks (Gow, Rogers, Jackson, & Schluter, 2008; Hatfield & Schluter, 1999; Schluter & McPhail, 1992; Walker, 1997; Willacker et al., 2010), Arctic charr (Brachmann, Parsons, Skulason, & Ferguson, 2021), minnows (Burress, Holcomb, Tan, & Armbruster, 2017), and carp (Hollingsworth, Simons, Fordyce, & Hulsey, 2013).

Benthic fishes live within rocks or shorelines and are thus in environments with variation in structure and water flow. Fishes in this habitat generally have a deeper, more stout body thought to be adaptive for increased maneuverability and body rotation (Webb, 1982, 1984). For instance, a deeper caudal peduncle and more posterior placement of the anal and dorsal fins enable benthic fishes to have fast propulsion as they move, change direction, and adjust to varied water flow patterns in their complex habitats (Koehl, 1984; Webb, 1982). This body variation can be coordinated with changes in head shape, namely a shorter head region (Cooper et al., 2010). On the other end of the axis, pelagic or limnetic fishes live in open waters of oceans or lakes, respectively. Here, we use “pelagic” as a more generalized form of this open water habitat and ecology. Pelagic fishes have repeatedly evolved a slender, more narrow fusiform body shape, many times with a larger head proportion (Cooper et al., 2010; S. T. Friedman et al., 2020; Schluter & McPhail, 1992). This narrower body shape is thought to enhance performance of sustained swimming (“cruising”) in open water while minimizing drag (Raffini, Schneider, Franchini, Kautt, & Meyer, 2020; Webb, 1982, 1984).

We examine the genetic basis of this benthic-pelagic body shape variation using a textbook example of adaptive radiation, cichlid fishes. Within cichlids, benthic-pelagic habitat divergence is the first stage of their adaptive radiation (Kocher, 2004; Streelman & Danley, 2003) and occurs repeatedly, independently, and convergently in three expansive radiations in East African Rift Lakes (Cooper et al., 2010; Hulsey, Holzman, & Meyer, 2018; Hulsey et al., 2013; Muschick et al., 2012; Ruber & Adams, 2001) as well as smaller radiations within New World crater lakes (Elmer, Kusche, Lehtonen, & Meyer, 2010; Franchini et al., 2014; Kusche et al., 2014). Notably, hybrids among difference cichlids species or even genera can be produced in the lab, enabling quantitative trait loci (QTL) mapping of phenotypic variation (Powder & Albertson, 2016).

We therefore capitalized on the extensive phenotypic variation and genetics of cichlids to investigate the genetic basis of body shape divergence associated with the benthic-pelagic axis, a major axis of fish diversification. First, we determined the genetics of body shape within each cross, using species at varying points along this morphological continuum. Then, we compare if the genetic intervals and mechanisms of divergence are similar between the two hybrid crosses. If so, this would indicate a shared molecular control whether a fish is evolving towards a narrower body shape as towards an extreme benthic species. Second, we assess the genetic architecture of body shape evolution in cichlids. We ask if there are many genetic loci, each with small effects, or few regions with large effects that regulate body shape. Body shape variation within stickleback fishes has been shown to be due to a combination of a few QTL with large effects and many QTL with small effects (Albert et al., 2008; Liu et al., 2014; Yang, Guo, Shikano, Liu, & Merila, 2016). Further, QTL for body shape and other morphological features in sticklebacks have been suggested to cluster on certain chromosomes, and these “supergene” regions can influence coordinated changes in phenotype (Albert et al., 2008; Liu et al., 2014; Miller et al., 2014). Finally, we ask if this common and predictable morphological trajectory in fishes has a predictable genetic basis by comparing QTL for body shape across benthic-pelagic divergence in multiple fish species. In other words, we determine if there is parallelism in the genetics of body shape variation to accompany convergent morphologies. To accomplish these goals, we utilized two hybrid crosses of Lake Malawi cichlids, quantifying a suite of linear and geometric measures of body shape. We use QTL mapping to identify the genetic bases of these traits, which we then compare among traits, between crosses, and to studies in other non-cichlid fishes. Together, these data provide insights into the genetic control of a major ecological and morphological divergence in animals.

## MATERIALS AND METHODS

### Animals and pedigree

All animal care was conducted under approved IACUC protocol 14-101-O at North Carolina State University. Four species of Lake Malawi cichlids were used: *Aulonocara koningsi, Metriaclima mbenjii, Labidochromis caeruleus*, and *Labeotropheus trewavasae*, hereafter referred to by the genus name. Though none of these species are truly pelagic, these fish serve as a phenotypic proxy for various points along the primary benthic-pelagic morphological and evolutionary axis in cichlid fishes (Figure 1)(Konings, 2016); it is unlikely that hybrid crosses would be viable between true pelagic and benthic cichlids. *Aulonocara* lives within an open, sandy region, away from the complex rocky habitat of the other species used here, and forages by cruising over the open sand. Similarly, the *Labidochromis* species studied here lives in and around rocky habitats, but is non-territorial and swims continuously and darts among rocks in search of invertebrate prey. *Metriaclima* is a generalist benthic fish, and *Labeotropheus* represents the extreme end of benthic morphology and behavior (Cooper et al., 2010; Konings, 2016). Two hybrid crosses of these species were generated. The first cross came from a single *Metriaclima* female crossed to two *Aulonocara* males; the inclusion of the second grandsire was inadvertent and resulted from an unexpected fertilization event in these species with external fertilization. The second cross came from a single *Labidochromis* female crossed to a single *Labeotropheus* male. Thus, both crosses feature a fish that dominantly “cruises,” a pelagic swimming tactic, versus a more benthic species. For each cross, a single F_1_ family was generated and subsequently incrossed to produce F_2_ hybrid mapping populations. F_2_ families were raised in density-controlled aquaria with standardized measured feedings until five months of age for analysis. Fish were anesthetized with buffered 100 mg/L MS-222 for all photographs. Whole fish photographs were taken including a color standard and scale bar under uniform lighting conditions in a lightbox with a mirrorless digital camera (Olympus). The sex of each animal was determined based on gonad dissection and these data were omitted if there was ambiguity in gonad phenotype.

**Figure 1.**
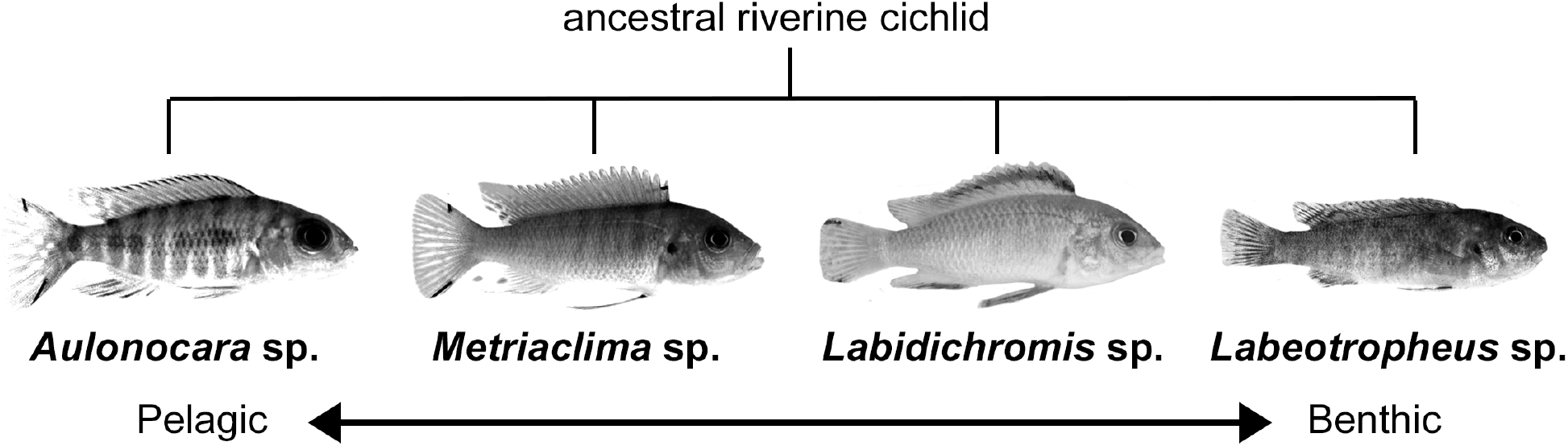
Parental species represent various points along the benthic-pelagic ecomorphological axis of Lake Malawi cichlids. Two F_2_ hybrid populations were generated. The benthic *Metriaclima* was crossed to *Aulonocara*, which lives in open sandy areas. Another cross was within benthic species, with *Labidochromis* crossed to the extreme benthic specialist *Labeotropheus*.

### Linear quantification of body shape variation

We quantified various measures of body shape in 10 individuals of each parental species, 491 *Metriaclima* x *Aulonocara* hybrids, and 447 *Labidochromis* x *Labeotropheus* hybrids. From photographs, we calculated series of linear distances (Figure 2b) using ImageJ software, including standard length (snout to caudal peduncle), head length (snout to opercle), body depth (anterior insertion of dorsal fin to insertion of pelvic fin), caudal peduncle width, distance between caudal peduncle and anal fin insertion, width of the anal fin base, distance between anal fin and pelvic fin, and pectoral fin width. ImageJ lengths in pixels were converted into centimeters using the scale bar included in each picture. To remove the effects of allometry on body measures, all measurements were converted into residual data by normalizing to the standard length, using a dataset including both parentals and their hybrids for a single cross. Analyses including linear normalization, ANOVAs, Tukey’s Honest Significant Difference post-hoc tests, and correlations were conducted in R.

**Figure 2.**
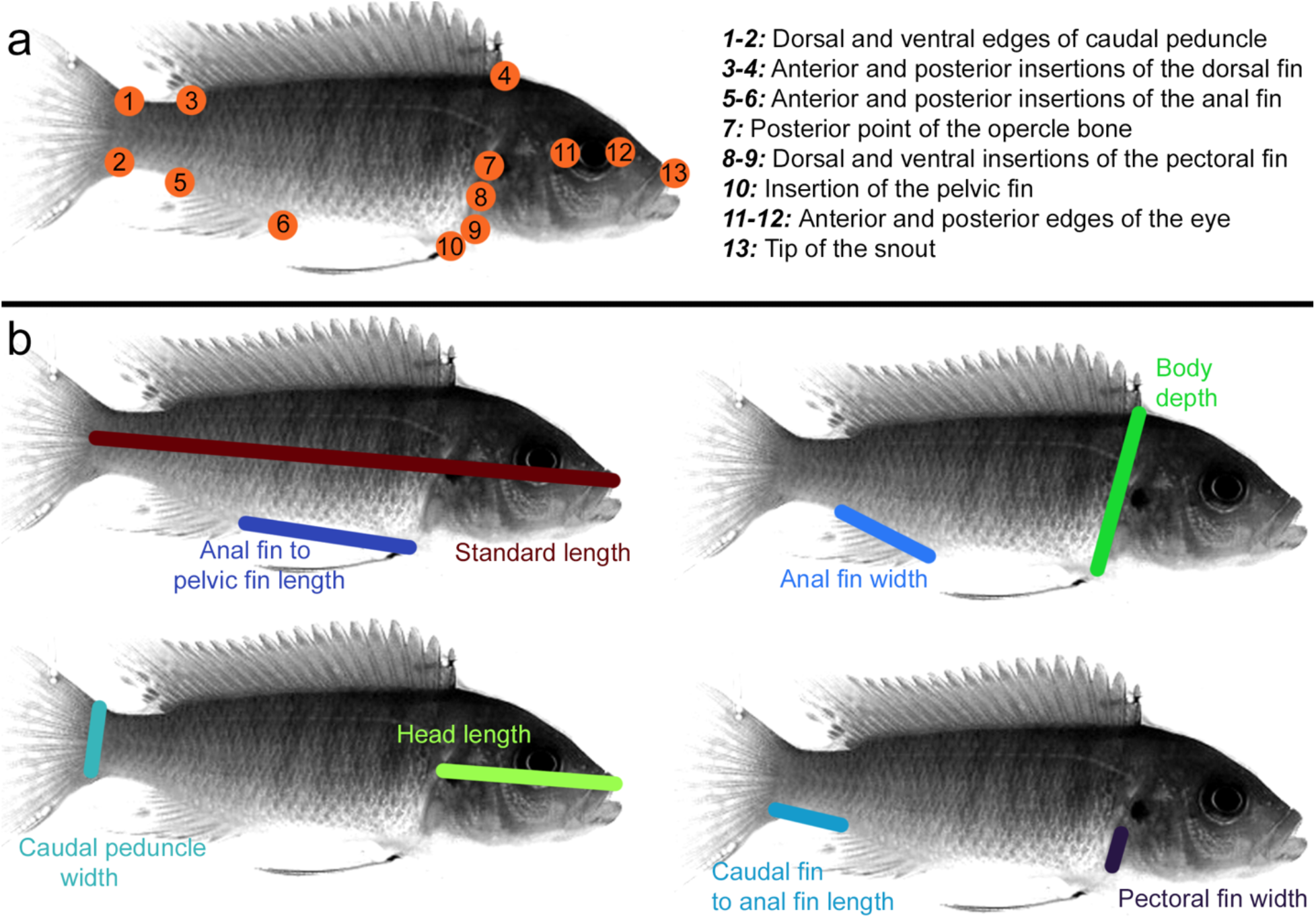
Measures used to assess body shape. **(a)** Geometric and (b) linear measures were used to assess body shape changes commonly seen along benthic-pelagic morphological divergence in fishes. Phenotypes are color-coded the same throughout figures.

### Geometric quantification of body shape variation

Body shape variation was also quantified using geometric morphometric shape analysis. Homologous anatomic landmarks were defined along the body primarily using fin insertion sites as well as head landmarks such as the snout, eye, and opercle (Figure 2a). Coordinate positions of all landmarks were collected from photos using the tpsDig2 software package (http://www.sbmorphometrics.org/). Landmark coordinates were extracted and inputted into the R package geomorph (Adams & Otarola-Castillo, 2013), which was used to conduct Procrustes superimposition of landmarks to remove variation due to size, rotation, and position of landmarks to leave variation only due to shape. The effects of allometry were removed with size correction and multiple regression of shape on standard length, using a data set including both parentals and their hybrids for a single cross.

### ddRAD-sequencing

Genomic DNA was extracted from caudal fin tissue using DNeasy Blood and Tissue kits (Qiagen). Indexed, double-digestion RADseq libraries were produced as previously described (Burford Reiskind et al., 2016) and sequenced on an Illumina HiSeq with 100 bp paired end reads (North Carolina State University Genomic Sciences Laboratory core facility). Raw sequencing data were demultiplexed, reads were truncated to 150 bp, and low-quality reads were excluded using the program process_radtags (Stacks, version 2). The demultiplexed and filtered reads were aligned to the *Maylandia zebra* UMD2a reference genome using BWA with the mem algorithm. We used the programs pstacks and cstacks (Stacks, version 1) to identify RAD markers in all the samples and create a catalogue of RAD markers present in both parents of the cross. The RAD markers of the progeny were subsequently matched against this catalog with the program sstacks (Stacks, version 1). Genotype calls for biallelic markers with alternative alleles between the parents of the cross (aa x bb markers) were generated with the program genotypes (Stacks, version 1), requiring a minimum stack depth of 3 in order to export a locus in a particular individual. For the *Metriaclima x Aulonocara* cross, markers were only used if both *Aulonocara* sires shared the same homozygous genotype.

### Linkage map

The genetic map was built on the R statistical platform (version 4.0.3) with the package R/qtl (Broman, 2009) and in-house scripts. RAD markers were sorted and binned in linkage groups according to their position in the *M. zebra* UMD2a reference genome. Markers located in linkage groups with more than 20% of missing data and markers located in unplaced scaffolds with more than 40% of missing data were filtered out from the dataset. A chi-square test was performed with the function geno.table() to detect markers with distorted segregation patterns; markers with a Bonferroni-corrected p-value < 0.01 were removed. The pairwise recombination frequencies among markers were calculated, and an initial map was estimated with the functions est.rf() and est.map(). Markers in linkage groups that did not show evidence of being misplaced were considered to be inflating the map and removed if they increased the size of the map by at least 6 centiMorgans (cM) and their flanking markers were less than 3Mb apart. Markers located in unplaced scaffolds were integrated into a given linkage group if they had recombination frequency values < 0.15 with at least five markers from that linkage group. The remaining markers located in unplaced scaffolds were removed from the map. Markers whose recombination frequency profile did not match their position in the genetic map, likely due to being located in structural variants or misassembled regions of the reference genome, were rearranged manually in order to minimize the number of crossovers. The function calc.errorlod() was used to detect genotyping errors; genotypes with a LOD score ≥ 3 were set as missing data. The map was pruned using a non-overlapping window algorithm that selected the marker in a given 2cM window with the least amount of missing data. The final map was estimated and the maximum likelihood estimate of the genotyping error rate (0.0001) was obtained with the function est.map(). The final genetic map for *Metriaclima* x *Aulonocara* hybrids included 22 linkage groups, 1267 total markers, 19-127 markers per linkage group, and was 1307.2 cM in total size. The final genetic map for *Labidochromis* x *Labeotropheus* hybrids included 22 linkage groups, 1180 total markers, 42-81 markers per linkage group, and was 1239.5 cM in total size.

### Quantitative trait loci (QTL) analysis

QTL mapping used multiple-QTL mapping (MQM) methods. Scripts are described in and available from (Powder, 2020), follow (Jansen, 1994), and use the R/qtl package (Arends, Prins, Jansen, & Broman, 2010; Broman, Wu, Sen, & Churchill, 2003). The process begins with a liberal scan for unlinked QTL using the onescan function in R/qtl (Broman, 2009). Putative QTL with a LOD approaching or above 3.0 were used to build a more rigorous statistical model. The MQM method use these putative loci as cofactors during a QTL scan, verified by backward elimination. The inclusion of cofactors in the final model provides more accurate detection of QTL and assessment of their effects (Jansen, 1994). Statistical significance of QTL was assessed using 1000 permutations. For QTL peaks meeting 5% (significance) or 10% (suggestive) level, 95% confidence intervals were determined using Bayes analysis. Scan details such as cofactors used and significance levels are reported in Table S1.

Markers are named based on contig and nucleotide positions in the *M. zebra* (zebra mbuna) reference genome, M_zebra_UMD2a assembly. Names, ID numbers, and start/stop positions of candidate genes within QTL intervals were extracted from the NCBI genome data viewer (https://www.ncbi.nlm.nih.gov/genome/gdv) gene track for *M. zebra* annotation release 104. If upper and lower limits of the 95% interval were markers that mapped to unplaced scaffolds, the closest marker that mapped to a placed scaffold was used instead. Gene names were compiled from the Database for Visualization and Integrated Discovery (DAVID) (Huang, Sherman, & Lempicki, 2009; Huang, Sherman, & Lempicki, 2009) using NCBI gene ID numbers as a query.

### Comparisons with other species

QTL intervals from other studies on body shape in different cichlid species (Franchini et al., 2014; Fruciano et al., 2016; Navon, Olearczyk, & Albertson, 2017), sticklebacks (Albert et al., 2008; G. L. Conte et al., 2015; Liu et al., 2014; Rogers et al., 2012; Yang et al., 2016), and carp (Laghari et al., 2014) were compiled, selecting for traits that were comparable with those studied here (i.e. similar measures to Figure 2). Chromosomal positions were converted to comparable intervals on the *M. zebra* (zebra mbuna) UMD2a genome assembly through a combination of manual BLAST of marker sequences and based on the locations of candidate genes within intervals. Specific phenotypes measured and details of conversion are detailed in Table S6.

## RESULTS

### Body shape variation

Body shape in our four parental species differs in measures such as body depth and relative head proportions, which are common changes across benthic-pelagic divergence in fishes. Parental species for both crosses are distinct in body shape, though not in all measures (Figures S1-S2 and Table S2). For both crosses, F_2_ hybrids are largely intermediate in phenotype to the two parental species, though phenotypic variation in hybrids is increased and can surpass the morphological range of parental species (Figures S1-S2). Thus, even for shapes with non-significant differences between parental species, QTL mapping can identify genetic loci that underlie shape variation (e.g. see pectoral fin width for both crosses in Figures 3-4 and Figures S1-S2).

**Figure 3.**
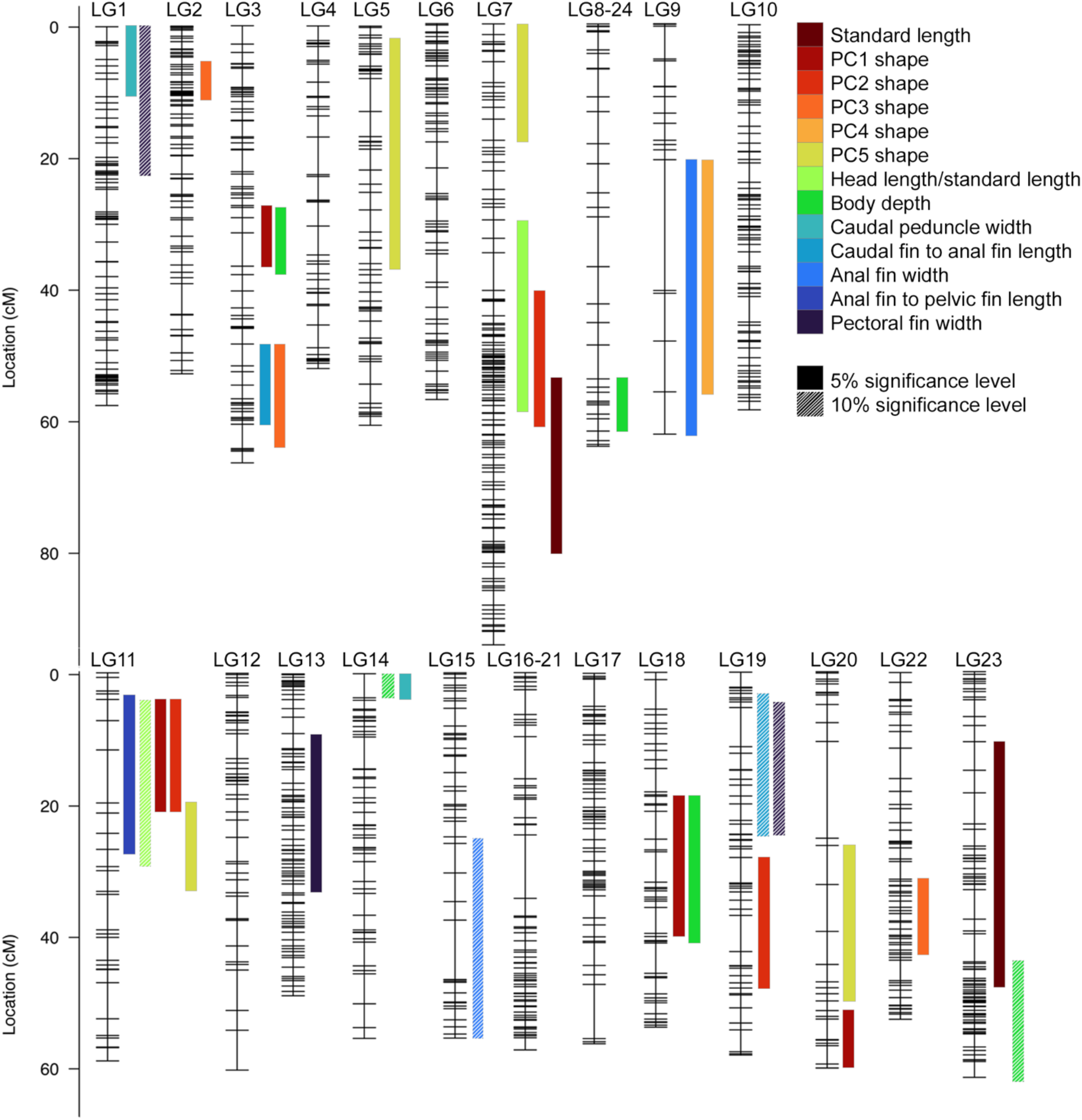
Quantitative trait loci (QTL) mapping identifies 34 intervals associated with changes in body shape between *Metriaclima* and *Aulonocara*. Each linkage group (LG, i.e. chromosome) is indicated with genetic marks noted by hash marks. The phenotype related to each QTL region is indicated by color. Solid bars are significant at the 5% genome-wide level, while those with white diagonals are suggestive, meeting the 10% genome-wide level. Bar widths indicate 95% confidence interval for the QTL, as calculated by Bayes analysis. Illustrations of phenotypes are in Figure 2, Figure S1, and Figure S3-4. QTL scans at the genome and linkage group level are in Figures S1 and S4. Details of the QTL scan including markers and physical locations defining each region are in Table S1.

**Figure 4.**
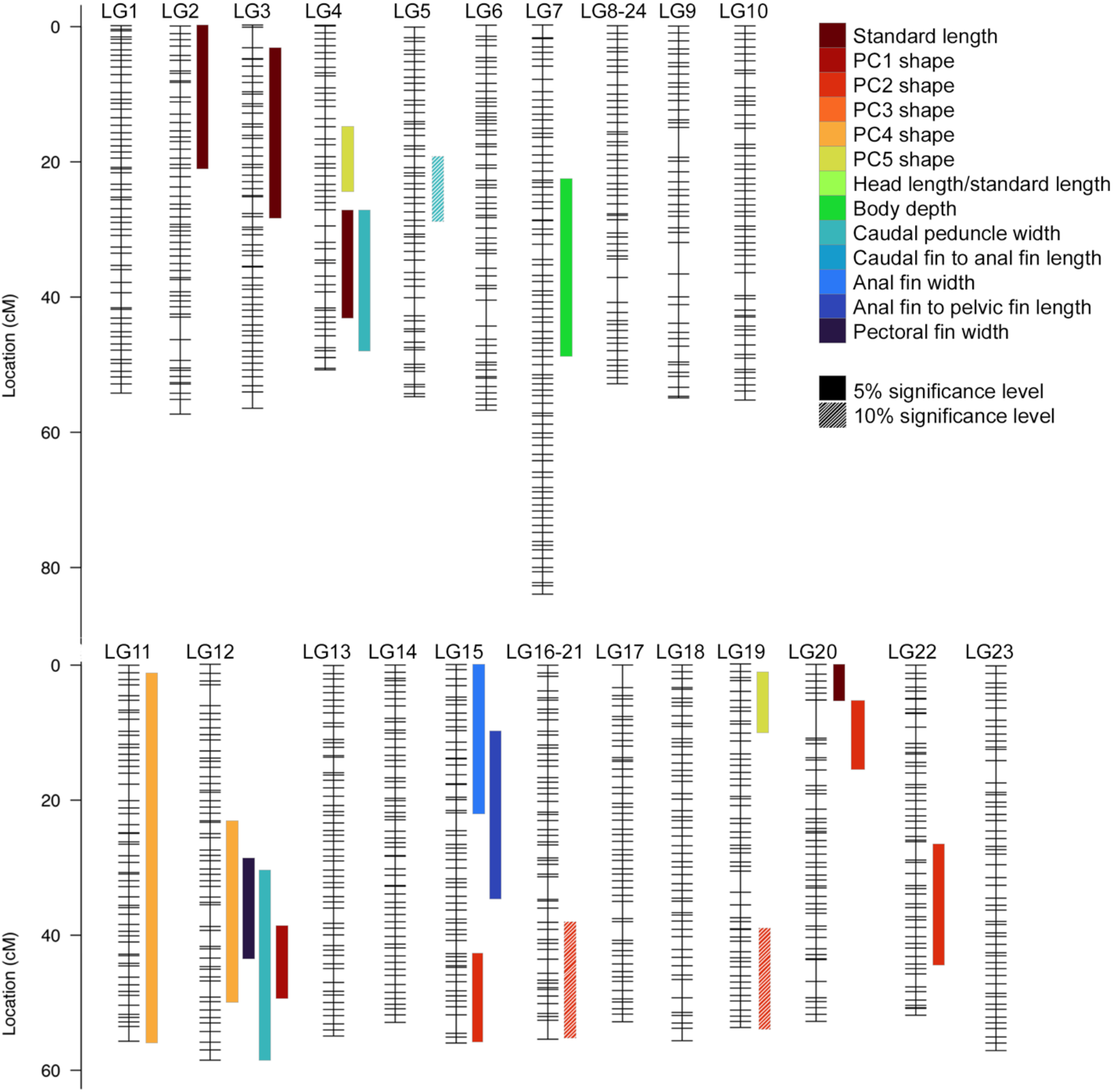
Quantitative trait loci (QTL) mapping identifies 21 intervals associated with changes in body shape between *Labidochromis* and *Labeotropheus*. Data is as presented in Figure 3.

Compared to *Metriaclima, Aulonocara* have significantly longer head proportions and a commensurate decrease in mid-body length as measured by the distance between the anal and pelvic fins (Figure S1 and Table S2). These differences are also supported by geometric morphometric shape analyses, which also identify coordinated changes in body shape. Together, the first five principal components describe 66.1% of total shape variation (TSV) in the bodies of *Metriaclima* sp., *Aulonocara* sp., and their F_2_ hybrids. Principal component 1 (PC1, 22.6% TSV) primarily describes variation in body depth, with positive scores associated with a deeper body commonly associated with benthic fishes, such as in cichlids (Elmer et al., 2010; Hulsey et al., 2018) and sticklebacks (Gow et al., 2008; Schluter & McPhail, 1992). PC2 scores (15.5% TSV) are associated with overall proportions of head versus flank in the body. In agreement with linear measurements, *Aulonocara* fishes are significantly associated with negative PC2 scores, which characterize a body with a larger head length, a posterior shift in pectoral and pelvic fins, and increased regions of the face anterior to the eye (Figure S1, Figure S3, and Table S2). Neither PC3 nor PC4 is significantly different between parentals. They are associated with head proportions, length of the caudal region, and the position of the anal fin (PC3, 11.7% TSV) and length of the caudal region and lengths of the anal fin (PC4, 8.4% TSV), respectively. Finally, PC5 (7.9% TSV) differs significantly between parentals, with *Aulonocara* fishes associated with a larger eye and more posteriorly positioned dorsal fin (Figure S1, Figure S3, and Table S2).

Despite both being rock-dwelling Lake Malawi cichlids that live in sympatry (Konings, 2016), *Labidochromis* and *Labeotropheus* parentals have more significant phenotypic differences than the *Aulonocara* and *Metriaclima* species pair (Figure 1 and Table S2). *Labeotropheus* species are highly derived, representing an extreme benthic morphology within Lake Malawi (Cooper et al., 2010), and have significantly shorter head regions, decreased body depth, decreased caudal peduncle width, decreased anal fin width, and increased length between the caudal peduncle and the anal fin compared to *Labidochromis* fish (Figure S2 and Table S2). As with the other cross, these trends can be identified in the geometric morphometric shape analysis, with the first five principal components explaining 64.8% of total shape variation in *Labidochromis* sp., *Labeotropheus* sp., and their F_2_ hybrids. Like the *Metriaclima* x *Aulonocara* cross, PC1 scores (17.7% TSV) primarily describe differences in body depth, and are not significant between parental species, nor between either parent and their F_2_ hybrids. However, all other PCs are significantly different between *Labeotropheus* and *Labidochromis* fishes. Positive PC2 scores (15.1% TSV) are associated with *Labidochromis*, an increased head length, increased body depth, and a longer anal fin (Figure S2, Figure S3, and Table S2). PC3 (12.9% TSV) characterizes *Labidochromis* fish as having a shorter distance between the caudal peduncle and anal fin, as well as a longer head region. PC4 (10.8% TSV) primarily describes fins on the ventral side of the fish, with *Labidochromis* and positive scores associated with a posterior shift of pectoral and pelvic fins and increased length of the anal fin. Finally, PC5 (8.3% TSV) illustrates that *Labidochromis* species have a deeper body shape and eyes positioned more ventrally on the head (Figure S2, Figure S3, and Table S2).

### Sexual dimorphism in body shape

Differences between males and females, termed sexual dimorphism, are incredibly common across species and often include differences in body size or shape (Badyaev, 2002; Fairbairn, Blackenhorn, & Szekely, 2007; Frayer & Wolpoff, 1985; Williams & Carroll, 2009). We therefore assessed differences between sexes in our two hybrid populations. Sex in Lake Malawi cichlids is genetically determined, with a diversity of sex determination loci identified (Gammerdinger & Kocher, 2018). Within the *Labidochromis* x *Labeotropheus* cross, sex is not associated with common sex determination loci, and of the 354 animals for which sex was called, 92.9% were male. In the *Metriaclima* x *Aulonocara* cross, sex is solely determined by a common XY system on LG7, which produces an even one-to-one sex ratio (Peterson, Cline, Moore, Roberts, & Roberts, 2017; Ser, Roberts, & Kocher, 2010). Thus, we only discuss sexual dimorphism based on 412 *Metriaclima* x *Aulonocara* hybrids (48.1% male), though see Table S2 for full ANOVA analyses in both crosses.

Overall size was the most significant body shape difference between males and females (standard length, Table S2). Within the linear measures of body shape, males had significantly larger head proportions, body depth, and width of the anal fin, which features egg spot pigmentation patterns used during mating (Table S2). This is supported by dimorphism in geometric morphometric shape analyses, with males being associated with increased body depth (positive PC1 score), a larger head (negative PC2 scores), and larger anal fin (negative PC4 scores) (Figure S3, Table S2).

### Genetic basis of body shape variation

To determine the genetic mechanisms that underlie variation in body shape, we genetically mapped five PC measures of complex shape changes (Figure S3) and eight linear measures (Figure 2). We did this in both the *Metriaclima* x *Aulonocara* cross and *Labidochromis* x *Labeotropheus* cross, which represent different benthic-pelagic divergences within Lake Malawi cichlids.

Thirty-four genetic intervals underlie the quantitative differences in body shape in *Metriaclima* x *Aulonocara* F_2_ hybrids. This includes 27 that reach 5% statistical significance at the genome-wide level and 7 that are suggestive, reaching 10% significance levels (Figure 3, Figure S1, Figure S4, Table S1). Each trait has 1-5 genetic intervals that influence phenotypic variation. QTL intervals are spread throughout the genome, with 16 of 22 linkage groups each containing 1-5 loci. QTL regions explain from 3.0-7.2% of the total variation in each trait (Figure S4), meaning that all body shapes analyzed are controlled by many genetic loci, each with a small effect.

Despite having more phenotypically distinct parentals (Table S2), fewer QTL were identified in *Labidochromis* x *Labeotropheus* F_2_ hybrids. A total of 18 significant and 3 suggestive QTL were identified, with 0-4 QTL per trait (Figure 4, Figure S2, Figure S5, Table S1). Like the other cross, QTL are spread throughout the genome, with 13 of 22 linkage groups containing 1-4 QTL each. Phenotypic traits are similarly complex in terms of genetics, with each QTL only accounting for 3.2-5.8% of shape variation (Figure S5). Even for the trait with the most variation explained (PC2 score in *Labidochromis* x *Labeotropheus*), only 21.0% of total shape is explained by the combined effects of 5 QTL (Figure S5, Table S1).

QTL between the two crosses are largely non-overlapping both generally and for specific traits (Figures 3-4, Figures S1-S5, Table S1). The linkage groups that contain “hotspots” with multiple QTL in one cross are largely absent from QTL in the other cross. For example, LGs 3, 7, and 11 contain 4, 4, and 5 QTL respectively in the *Metriaclima* x *Aulonocara* cross; each of these LGs only contain one QTL in the *Labidochromis* x *Labeotropheus* cross. Alternately, the linkage groups with the most QTL in the *Labidochromis* x *Labeotropheus* cross (LG4 with 3 QTL and LG12 with 4 QTL), do not contain any QTL in the *Metriaclima* x *Aulonocara* cross.

This lack of common genetic signals is also the case when looking at individual traits. By and large, the same trait is explained by distinct genetic intervals in each cross (Figures 3-4). In both crosses, LG15 contains a QTL for anal fin width. However, the 95% confidence intervals do not overlap, residing 23 Mb away from each other (Figures 3-4, Table S1). Thus, two distinct genetic intervals control anal fin width in these crosses, but both happen to occur on the same linkage group. There is also an overlap in QTL explaining PC2 scores in each cross, found on LG19. The significant QTL in the *Metriaclima* x *Aulonocara* cross overlaps with a suggestive QTL in the *Labidochromis* x *Labeotropheus* cross by 2.66 Mb (Figures 3-4, Table S1). It is notable that the shapes explained by PC2 in each cross are similar, but with differences. In both cases, this variation includes differences in head length. However, PC2 in the *Labidochromis* x *Labeotropheus* cross also includes coordinated changes in body depth and anal fin length that do not occur in the other cross (Figure S3).

However, there are some overlapping QTL within each cross, indicating that certain genomic intervals may have pleiotropic effects on body shape (Figures 3-4, Table S1). First, three QTL from the *Metriaclima* x *Aulonocara* cross overlap from 56.3-58.0 cM on LG7, contributing to variation in standard length, head proportion, and PC2 shape that also describes head proportions (Figure 3). Notably, all three traits are sexually dimorphic (Table S2), and the Y sex determining locus is nearby at 61.4-62.9 cM (Peterson et al., 2017). Second, a QTL hotspot in the *Metriaclima* x *Aulonocara* cross occurs from 3.2-33.3 cM on LG11. Five QTL reside in this region and explain disparate variation in body shape, including head proportions (linear measure as well as PC2 score), the distance between the anal and pelvic fins, body depth (as PC1 score), and variation in eye position and dorsal fin positioning (as PC5 score). This “hotspot” may be explained by a large, previously-characterized chromosomal inversion on LG11 between *Metriaclima* sp. genomes and *Aulonocara* sp. genomes (M. A. Conte et al., 2019). Finally, a QTL hotspot in the *Labidochromis* x *Labeotropheus* cross occurs from 23.5-58.8 on LG12. This region underlies phenotypic variation in four distinct body shapes: pectoral fin width, caudal peduncle width, body depth (as PC1 score), and positioning of the pectoral and pelvic fins (as PC4 score).This region of LG12 is also notable for a series of structural differences and changes in recombination rates among cichlid species (M. A. Conte et al., 2019). Thus, at each interval where multiple QTL overlap, interspecific variation in structural features of the genome are implicated.

There are no clear trends when looking at allelic effects on phenotypes in either cross (Figures S4-S5, Table S1). For example, standard length in the *Labidochromis* x *Labeotropheus* cross is not significantly different between species, is controlled by four QTL, and at all four of these loci the derived *Labeotropheus* allele increases size, through a mixture of additive, overdominant, and underdominant inheritance (Figure S5a). However, for PC2 shape in the same cross, which is significantly higher in *Labidochromis*, the allele from this species only increases trait values for three of the five underlying QTL (Figure S5c).

Additional work will be needed to further clarify the molecular mechanisms through which phenotypic variation is generated. While some intervals have relatively few candidate genes (e.g. QTL for body depth on LG14 with only 11 genes), most of our intervals contain hundreds of genes (Tables S4-S5). While it is too early to speculate on the effects of specific candidate genes in these intervals, we looked for overlap between our body QTL and three genes (*bmp4, lbh*, and *ptch1*) previously implicated in trophic adaptations that commonly co-occur with body shape variation in cichlids (Cooper et al., 2010). *Bmp4* and *lbh* reside close together on LG19 and regulate mandible length (Albertson, Streelman, Kocher, & Yelick, 2005) and early facial patterning (Powder, Cousin, McLinden, & Albertson, 2014), respectively. These genes co-localize with a QTL for PC2 shape in the *Metriaclima* x *Aulonocara* cross, which largely describes changes in head length. *Ptch1* variation produces alternate shapes in the lower jaw that represent a tradeoff between two feeding mechanisms: suction feeding associated with more pelagic species and biting that is common within specialized benthic species (Roberts, Hu, Albertson, & Kocher, 2011). *Ptch1* co-localizes with the LG12 QTL hotspot in the *Labidochromis* x *Labeotropheus* on LG12. Though this region is also associated with altered patterns of recombination, this leaves open the possibility that a gene such as *ptch1* may have pleiotropic effects on multiple adaptations associated with benthic-pelagic divergence.

### Modularity of body shape variation

The presence of genetic hotspots linked to multiple phenotypes suggests that a single locus may have pleiotropic effects on multiple aspects of body shape variation. To directly address the relationships among our measured phenotypes, we assessed correlations between all pairs of traits. For both crosses, standard length was positively and strongly associated with linear measures (e.g. r^2^ values from 0.846-0.963 in the *Metriaclima* x *Aulonocara* cross, Table S3). Note that correlations between standard length and PC scores were conducted following a size correction. After removing the effects of size, residuals for each measure were not strongly correlated with each other (r^2^ values from -0.562 to 0.713 with a mean of 0.005 in the *Metriaclima* x *Aulonocara* cross, and r^2^ values from -0.412 to 0.423 with a mean of 0.017 in the *Labidochromis* x *Labeotropheus* cross) (Table S3). This lack of correlation suggests that despite some traits having overlapping genetic influences (i.e. QTL), the phenotypic patterns generated are distinct. While individual linear measures did not correlate well with PC scores in the *Labidochromis* x *Labeotropheus* cross, PC scores seemed to be more strongly driven by single phenotypic metrics in the *Metriaclima* x *Aulonocara* cross. Specifically, PC1 score correlated with body depth (0.713), PC2 score correlated with head proportions (−0.562), PC3 score correlated with the distance between the caudal peduncle and anal fin (0.612), and PC4 score correlated with anal fin width (−0.540) (Table S3). While these are weak correlations, this suggests that the shapes described by geometric morphometrics in the *Metriaclima* x *Aulonocara* may be simpler than in the other cross, and variation between these pelagic and benthic species is more modular.

## DISCUSSION

### Evolution of an ecologically important trait

We sought to understand the genetics of body shape along the benthic-pelagic axis using multiple hybrid crosses of Lake Malawi cichlids and a series of both linear and geometric shape measurements. We find numerous differences in body shape including body depth, head proportions, caudal peduncle depth, and shifting of fin insertions that are likely to have functional consequences for swimming movements. This series of changes in the body plan seen in the cichlids used here mirrors variation commonly seen along the benthic-pelagic axis in fishes. Further, divergence of body shape between sympatric *Labidochromis* and *Labeotropheus* emphasize that while these fish encounter similar functional challenges of living in a complex benthic habitat, their distinct trophic niches are key factors driving variation in body shape. Specifically, the insectivore *Labidochromis* has larger head proportions, increased body depth, a deeper caudal peduncle, a shorter caudal region, and a longer anal fin than the algae-scraping *Labeotropheus* (Figure S2, Table S2) (Konings, 2016). These adaptations in *Labidochromis* are similar to fishes that use quick bursts of speed with abrupt shifts in direction (Meyers & Belk, 2014; Webb, 1984), and may reflect an ability of *Labidochromis* to quickly maneuver and pursue insect prey while *Labeotropheus* feed by hovering and holding steady within water flows of the lake to graze on algae.

We then used quantitative loci mapping to identify the genetic basis of these differences in body shape. We predicted that the same genetic intervals would influence body shape traits in both crosses. In other words, producing body shapes associated with a benthic environment would be the same regardless of relative position along the benthic-pelagic axis. However, we found that QTL that control body shape variation between the open water *Aulonocara* and benthic *Metriaclima* fishes are distinct from those that influence variation between a benthic generalist like *Labidochromis* and the benthic specialist *Labeotropheus* (Figures 3-4, Table S1). Coupled with modest effects of each QTL (3.0-7.2% variation explained by each), this emphasizes that body shape is a complex, polygenic trait and similar morphologies can be produced by multiple mechanisms (a many-to-one relationship).

### Distinct genetic signals regulate body shape across fishes

Body elongation in fishes along the benthic-pelagic axis occurs repeatedly and widely across fish phylogeny, independent of time (modern versus historic) and environment (marine versus freshwater) (Burns & Sidlauskas, 2018; S. T. Friedman et al., 2020; Gow et al., 2008; Hulsey et al., 2013; Muschick et al., 2012; Ribeiro et al., 2018; Robinson & Wilson, 1994). In “replaying life’s tape” (Gould, 1989), it is clear that fishes have converged on similar body shapes based on ecological selection. However, it is unknown if fish species use similar genetic mechanisms to achieve these predictable morphologies. Towards this goal, we compared the QTL in this study with QTL identified in additional Lake Malawi species (Navon et al., 2017), crater cichlids from Central America (Franchini et al., 2014), parallel radiations of sticklebacks (Albert et al., 2008; G. L. Conte et al., 2015; Rogers et al., 2012; Yang et al., 2016), and carp (Laghari et al., 2014) (Figure 5, Table S6). This comes with caveats such as missing data due to unclear orthologous regions in the *M. zebra* genome used here, the inclusion of only some fish species, and similar, yet not identical measures in other studies. Despite this, our comparative approach can still identify if these parallel benthic-pelagic divergences and changes in body shape are due to common genetic mechanisms across multiple fish clades.

**Figure 5.**
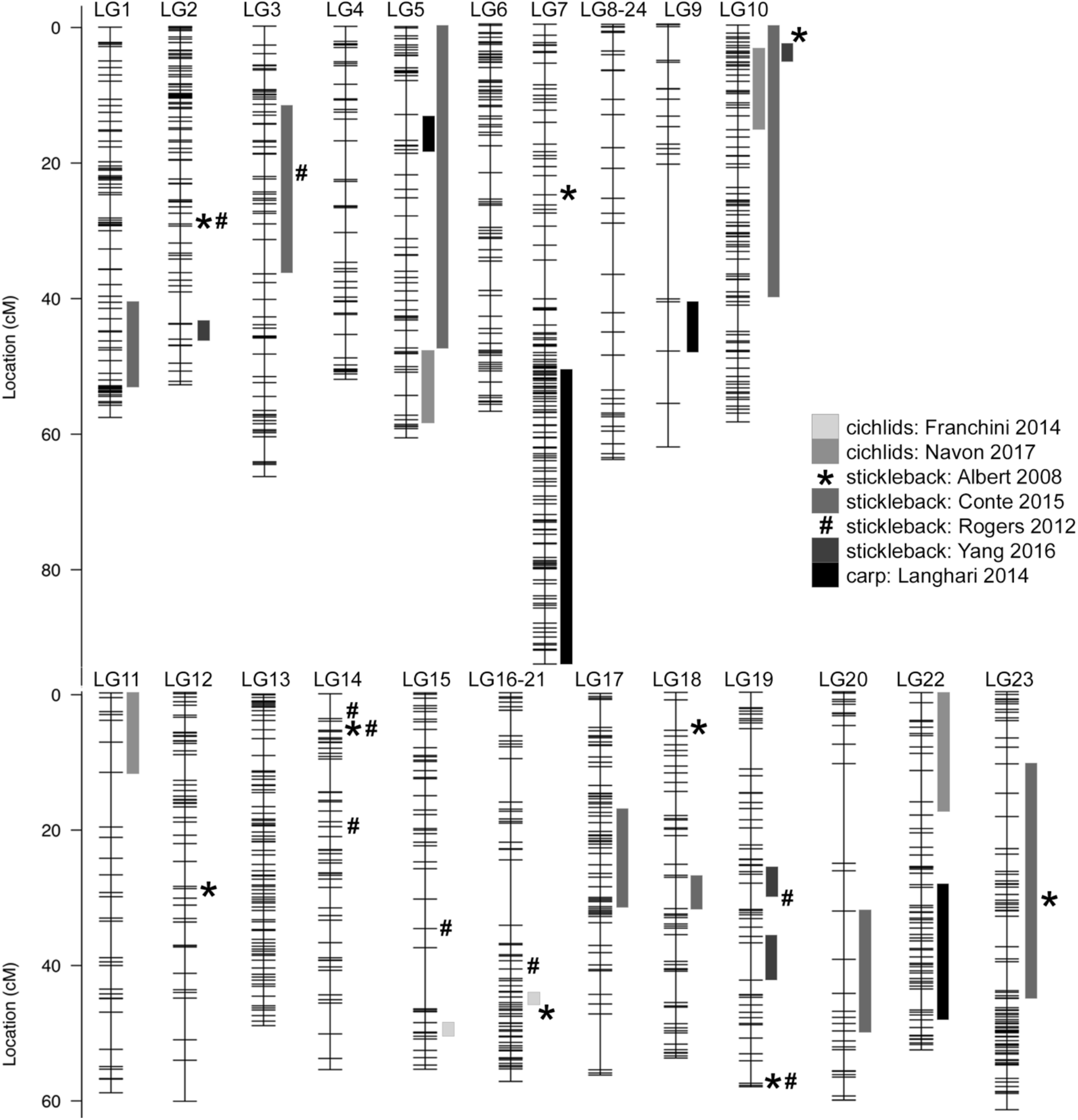
Comparison with QTL intervals previously identified in other cichlids, stickleback, and carp reveals little overlap with QTL determined in this study. Reported QTL from the indicated reference were converted to physical locations in the *M. zebra* genome UMD2a assembly and mapped onto the *Metriaclima* x *Aulonocara* genetic map. Bar widths indicate the 95% confidence interval for the QTL as calculated by each study. Those studies that reported QTL peak positions, but not confidence intervals, are indicated by * and # symbols. Details of phenotypes associated with each interval, physical locations, and methods of converting to UMD2a genome positions are detailed in Table S6.

Overall, we find relatively minimal overlaps in QTL among our study and QTL analysis of body shape in other fishes (Figures 3-5). For instance, while LG10 has been implicated in three of seven previous analyses (Figure 5), neither of our crosses had a QTL on this chromosome. Likewise, regions that have multiple QTL in this work (e.g. LG11 for the *Metriaclima* x *Aulonocara* cross or LG12 for the *Labidochromis* x *Labeotropheus* cross) have not been identified in previous analyses. Surprisingly, this trend even holds true for a cross within Lake Malawi (Navon et al., 2017), including another *Labeotropheus* species that is a close sister taxon to the one used here. Of four QTL previously associated with body shape variation in the extreme benthic *Labeotropheus* species, three (LG5, LG10, and LG22) show no overlap with QTL from either of our crosses. The fourth QTL lies on LG11, and overlaps with four QTL from the *Metriaclima* x *Aulonocara* cross, but not from the more similar *Labidochromis* x *Labeotropheus* cross. When some overlaps do occur, the traits do not always demonstrate a clear trend. For example, the LG19 region from 25-50cM shows an overlap of four QTL. While PC2 shape from both the *Metriaclima* x *Aulonocara* and *Labidochromis* x *Labeotropheus* crosses describes differences in head proportions, QTL in sticklebacks includes those for unrelated traits like caudal peduncle length and body depth (Figure 5, Table S6).

### Impacts of evolutionary history on genetic architecture and modularity

Much of benthic-pelagic body shape variation in stickleback fish is caused by few genes with large effects (10-20% variation explained each) and clustered “supergene” regions that influence multiple phenotypes associated with benthic ecologies (Albert et al., 2008; Liu et al., 2014; Miller et al., 2014; Yang et al., 2016). In contrast to this, we find that body shape variation in Lake Malawi cichlids is due to many genes, each with a small effect (Figures S4-S5, Table S1 and Tables S4-5). Additionally, we find that QTL are largely spread throughout the genome (Figures 3-4), and there are minimal correlations between measurements (Table S3). The largely modular basis of body shape variation in cichlids has implications for evolutionary potential (i.e. evolvability) and phenotypic variation (Melo, Porto, Cheverud, & Marroig, 2016; Pigliucci & Muller, 2010; Wagner, Pavlicev, & Cheverud, 2007). Namely, as distinct phenotypes are largely controlled by independent genetic intervals in cichlids, this allows independent evolution of distinct morphologies, each of which could be subject to different patterns of selection.

While most of the genetic correlations identified show independent segregation and phenotypic impacts, there are exceptions. One is found at the sex determination locus on LG7 in the *Metriaclima* x *Aulonocara* cross, where the multiple traits mapped to this region are also sexually dimorphic. Our study is unable to disentangle whether the LG7 associated trait variation is sex limited (i.e. modulated by sex-specific physiology during development, where sex is correlated to LG7 genotype), or results from allelic variation that is in linkage with the sex determination locus. If future studies indicate the latter, this would support the hypothesis that sexual dimorphism in body morphology evolves from accumulation of sexually antagonistic alleles at sex determination loci, as has been suggested for sexual dimorphism in pigmentation (Albertson et al., 2014; Roberts, Ser, & Kocher, 2009). Two other hotspots containing multiple trait QTL were found, one on LG11 in the *Metriaclima* x *Aulonocara* cross, and the other on LG12 in the *Labidochromis x Labeotropheus* cross. There is clear evidence for an interspecific inversion at LG11 between *Metriaclima* and *Aulonocara*, and recombination patterns among different hybrid crosses suggest that LG12 has significant variation in structure among Lake Malawi cichlids (M. A. Conte et al., 2019). Inversions and other structural variants can strongly suppress recombination, and this could support the evolutionary accumulation of complimentary adaptive alleles at multiple loci within broad haplotypes. Similar roles for inversions have been suggested in the parallel adaptation of sticklebacks (Jones et al., 2012), including the predictable fixation of certain inversion haplotypes within freshwater populations (Roesti, Kueng, Moser, & Berner, 2015). Within the Lake Malawi radiation, where occasional interspecific hybridization occurs and likely supports evolution, inversions may preserve combinations of alleles that drive multiple, distinct traits in the same direction along the benthic-pelagic axis, while avoiding discordant phenotypes. Our observations indicate a need for additional work dissecting genetic variation at structural variants. This could determine whether multiple genes are involved, or whether the hotspots represent the pleiotropic effects of a single gene that happens to lie within a structural variant.

This different genetic architecture and pattern of modularity within stickleback and cichlid fishes is likely due to their different evolutionary histories and ancestral states. Specifically, ancestral sticklebacks are marine, pelagic morphs that have a strong selective pressure towards a freshwater, benthic form when migrating into small lakes created by glacial retreat (Schluter & McPhail, 1992; Walker, 1997). On the other hand, cichlids evolved from a small group of generalist, riverine species (Malinsky et al., 2018) in sympatry towards multiple adaptive peaks. While benthic-pelagic habitat divergence is the first stage of the cichlid radiation and influences patterns of speciation, this is only one of many selective pressures (Kocher, 2004; Streelman & Danley, 2003). Further, the cichlid “hybrid swarm” has extensive shared genetic variation and ongoing hybridization among fish at varying points along the benthic-pelagic axis, which further influences the genetic architecture, modularity, and evolutionary potential of morphological variation in cichlids (Brawand et al., 2014; Malinsky et al., 2018).

## CONCLUSIONS

Body elongation is common across animals. In fishes, the benthic-pelagic ecomorphological axis is a major source of phenotypic variation, encompassing a suite of body shape phenotypes. We show here that this convergent phenotype is most likely due to distinct molecular signals in different fish clades or even at different points along a morphological continuum in a single radiation. Through comparison of genetic mapping results in two hybrid crosses, we show here that even the closely related cichlid species examined have distinct genetic architectures for this convergent trait. The genetic loci we identify here additionally serve as candidates to understand the molecular origins of an ecologically relevant trait, body shape variation.

## ACKNOWLEDGEMENTS

We thank Dr. Samantha Price for useful discussion and Dr. Daniela Almeida for useful comments on initial manuscripts. This work was supported by NSF CAREER #1942178 (KEP), NIH P20GM121342 (KEP), and NIH R15DE029945 (KEP), NSF IOS-1456765 (RBR), and an Arnold and Mabel Beckman Institute Young Investigator Award (RBR).

## DATA ACCESSIBILITY

Raw data files are available at Dryad [link to be provided prior to publication]. These files include phenotypic measures, TPS files for geometric morphometric analysis, and genotypes used for quantitative trait loci mapping.

## AUTHOR CONTRIBUTIONS

KEP and RBR designed the research. ACB, ECM, PJC, and NBR performed animal husbandry, photography, and collections. NBR prepared sequencing libraries. KEP, DM, and MJ performed phenotypic measurements. KEP, LD, ECM, ACB, and RBR analyzed data. KEP, LD, and AAB wrote the paper with edits from all authors.

