## Supplementary Data for "Genetic basis of body shape variation along the benthic-pelagic axis in cichlid fishes"

Supplemental table legends and figures with legends for DeLorenzo et al.


1. Legends for Supplemental tables, provided as separate .xlsx files (pg. 1-2)
    a. Table S1: Details of quantitative trait loci (QTL) identified in both crosses
    b. Table S2: Statistical significance of phenotypic measures
    c. Table S3: Correlations between all pairs of phenotypic traits
    d. Table S4: Candidate genes for body shape QTL identified in the *Metriaclima* x *Aulonocara* cross
    e. Table S5: Candidate genes for body shape QTL identified in the *Labidochromis* x *Labeotropheus* cross
    f. Table S6: Comparison to body shape QTL identified in other studies
2. Figure S1: Phenotypic measures and QTL scans within the *Metriaclima* x *Aulonocara* cross (pg. 3-6)
3. Figure S2: Phenotypic measures and QTL scans within the *Labidochromis* x *Labeotropheus* cross (pg. 7-10)
4. Figure S3: Shapes indicated by PC scores following geometric morphometric analysis (pg. 11)
5. Figure S4: QTL scans and allelic effects for significant QTL in *Metriaclima* x *Aulonocara* cross (pg. 12-25)
6. Figure S5: QTL scans and allelic effects for significant QTL in *Labidochromis* x *Labeotropheus* cross (pg. 26-37)


**Table S1. Details of quantitative trait loci (QTL) identified in both crosses.** For each QTL, we include cofactors use to generate models as well as markers and physical positions for the peak of the QTL and the 95% confidence interval. Marker names include the physical location on the linkage group, with names referring to the contig and nucleotide position in the *M. zebra* UMD2a assembly. LOD values, percent phenotypic variance explained by that QTL, allelic effects, and additive, dominance, and heritability calculations are for the peak marker in the QTL. QTL listed in gray are suggestive at the 10% significance level, while those in black meet 5% genome-wide significance based on values indicated.

**Table S2. Statistical significance of phenotypic measures.** Effects of size (standard length) and sex were assessed by ANOVA analysis. PC scores from geometric morphometric analysis were size corrected prior to this ANOVA. Significance between parental and hybrid groups were assessed for size-corrected, residual values followed by Tukeys HSD.

**Table S3. Correlations between all pairs of phenotypic traits**. Calculations are for $F_2$ hybrids animals. Except for correlations with standard length, all measures are size corrected. PC scores from geometric morphometric analysis were size corrected prior to assessing correlation with standard length. Values towards the top and right are for the *Metriaclima* x *Aulonocara* cross and those towards the bottom and left are for the *Labidochromis* x *Labeotropheus* cross. Values that are >0.85 or <-0.85 are in bold.

**Table S4. Candidate genes for body shape QTL identified in the *Metriaclima* x *Aulonocara* cross.** For the indicated 95% confidence interval, we include candidate gene names in the *M. zebra* annotation release 104, NCBI gene ID number, and physical positions in the *M. zebra* genome UMD2a assembly.

**Table S5. Candidate genes for body shape QTL identified in the *Labidochromis* x *Labeotropheus* cross.** Data is as presented in Table S4.

**Table S6. Comparison to body shape QTL identified in other studies.** Details of phenotype measured and reported QTL interval or marker from the indicated reference. Method of identifying the orthologous region in *M. zebra* UMD2a assembly is described and presented with physical and genetic locations visualized in Figure 5.

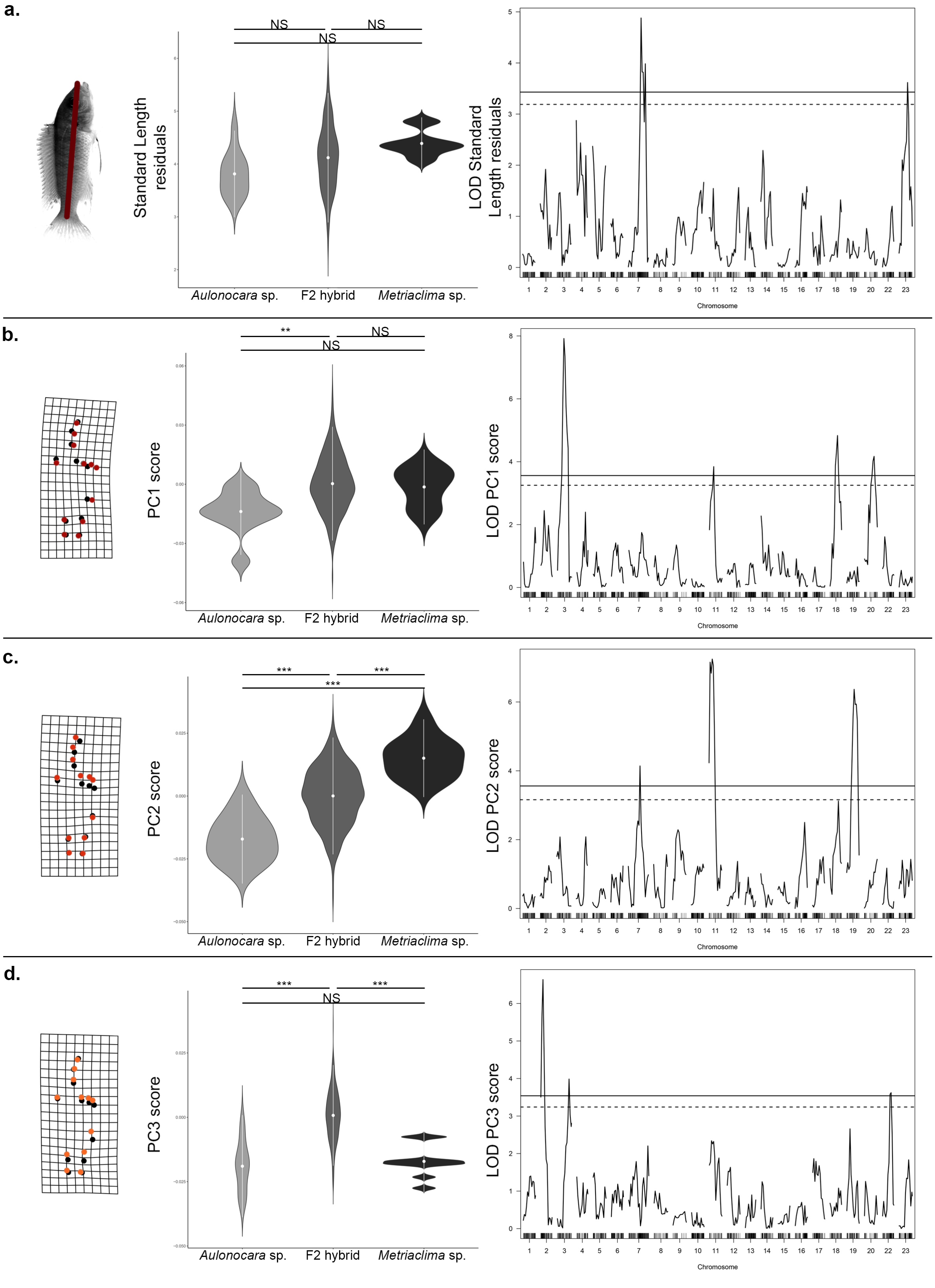
a.
NS
NS
NS
Standard Length residuals
Aulonocara sp.
F2 hybrid
Metriaclima sp.
LOD Standard Length residuals
Chromosome
b.
**
NS
NS
PC1 score
Aulonocara sp.
F2 hybrid
Metriaclima sp.
LOD PC1 score
Chromosome
c.
***
***
***
PC2 score
Aulonocara sp.
F2 hybrid
Metriaclima sp.
LOD PC2 score
Chromosome
d.
***
***
NS
PC3 score
Aulonocara sp.
F2 hybrid
Metriaclima sp.
LOD PC3 score
Chromosome

**e.**

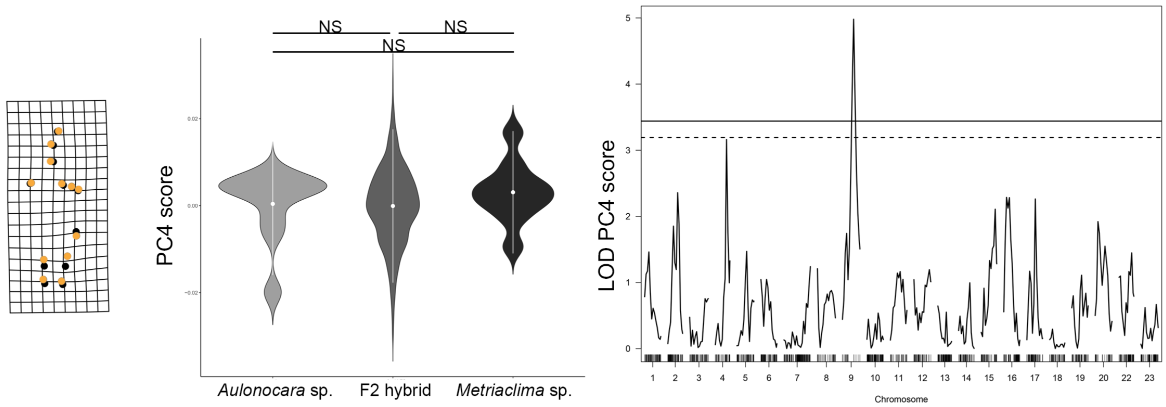
NS
NS
NS
PC4 score
Aulonocara sp.
F2 hybrid
Metriaclima sp.
LOD PC4 score
Chromosome

**f.**

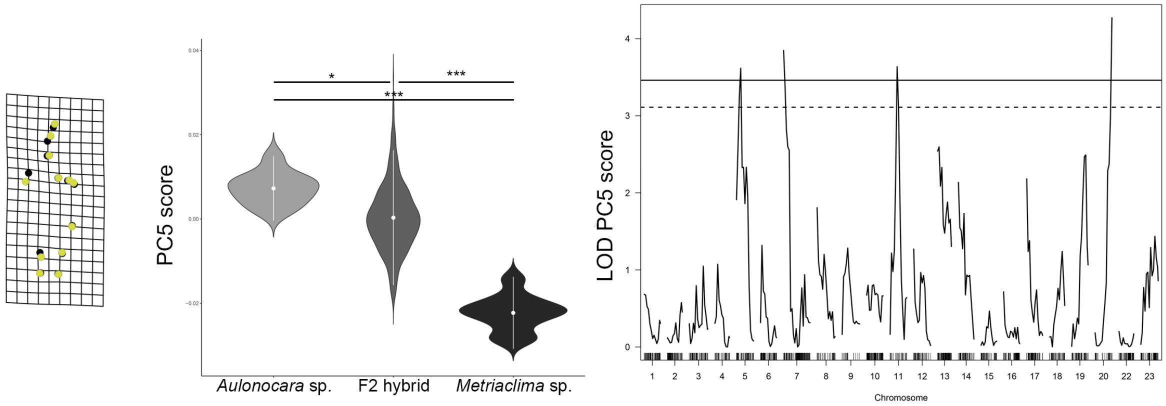
*
***
***
PC5 score
Aulonocara sp.
F2 hybrid
Metriaclima sp.
LOD PC5 score
Chromosome

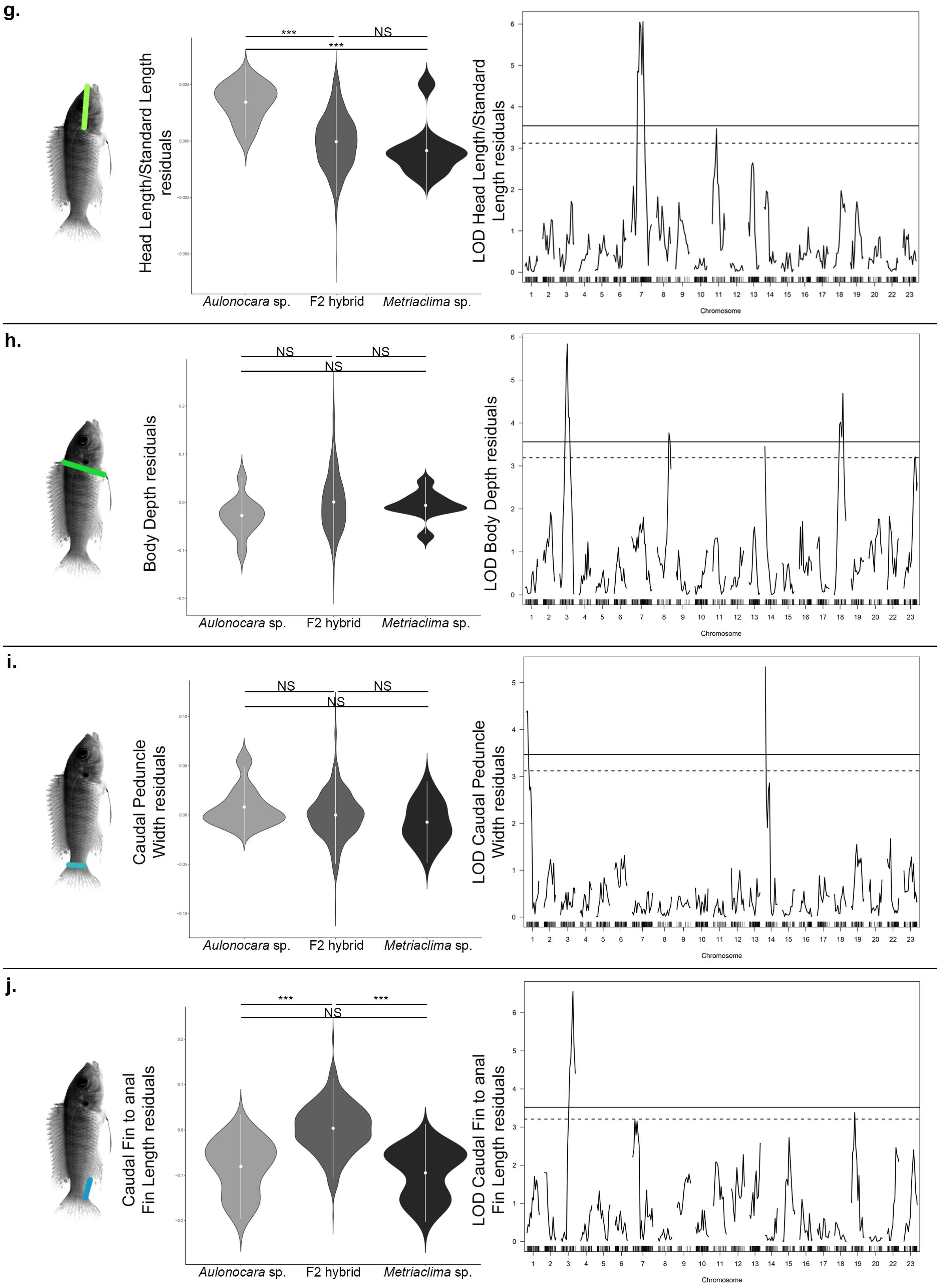
g.
***
NS
***
Head Length/Standard Length residuals
LOD Head Length/Standard Length residuals
Aulonocara sp.
F2 hybrid
Metriaclima sp.
Chromosome
h.
NS
NS
NS
Body Depth residuals
LOD Body Depth residuals
Aulonocara sp.
F2 hybrid
Metriaclima sp.
Chromosome
i.
NS
NS
NS
Caudal Peduncle Width residuals
LOD Caudal Peduncle Width residuals
Aulonocara sp.
F2 hybrid
Metriaclima sp.
Chromosome
j.
***
***
NS
Caudal Fin to anal Fin Length residuals
LOD Caudal Fin to anal Fin Length residuals
Aulonocara sp.
F2 hybrid
Metriaclima sp.
Chromosome

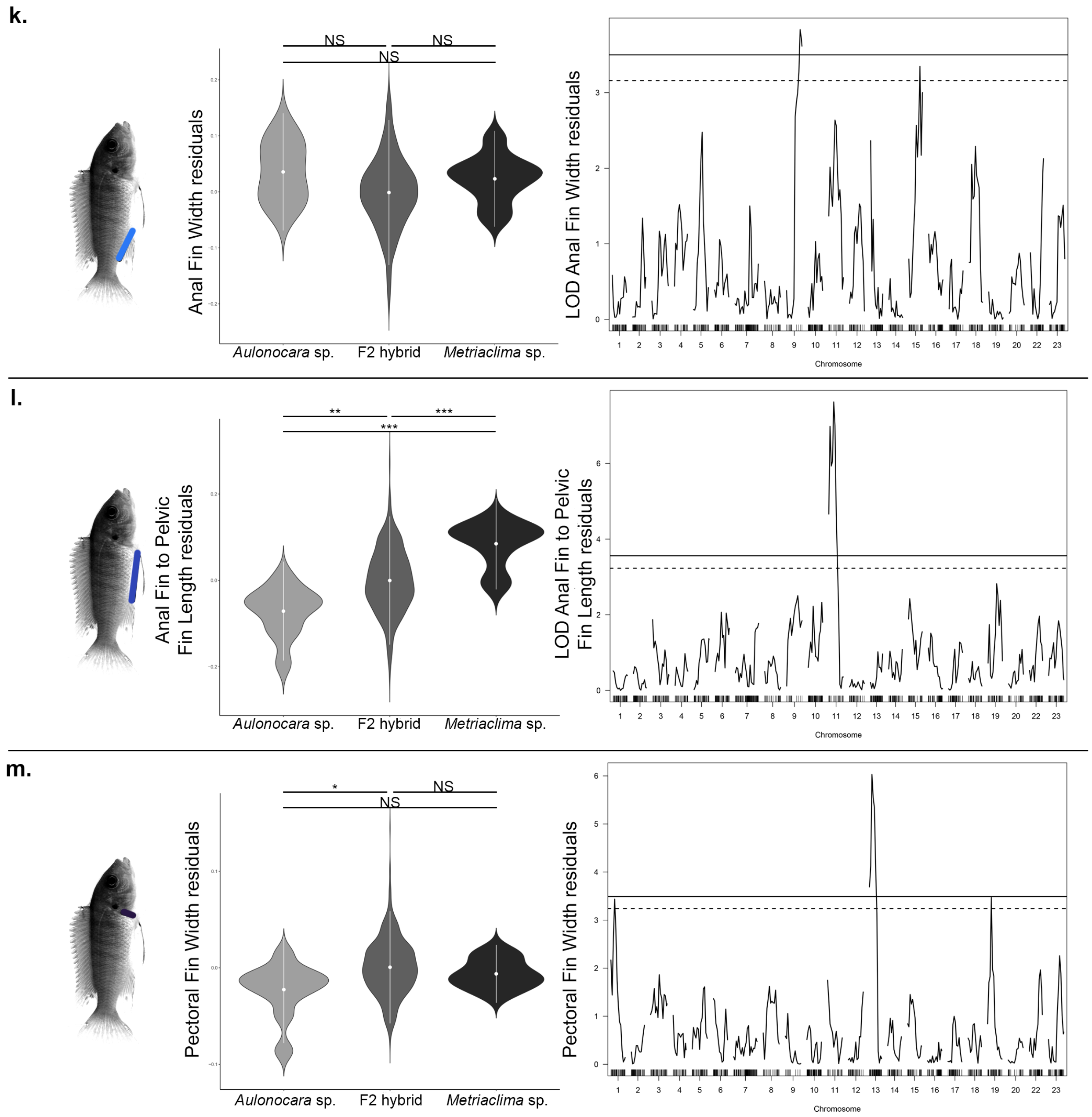


**Figure S1. Phenotypic measures and QTL scans within the *Metriaclima* x *Aulonocara* cross.** Phenotypic measures are indicated by illustration with colors matching Figures 2 and S3 and include (a) standard length, (b) PC1 shape, (c) PC2 shape, (d) PC3 shape, (e) PC4 shape, (f) PC5 shape, (g) head proportion, measured as head length/standard length, (h) body depth, (i) caudal peduncle width, (j) length between the caudal and anal fins, (k) anal fin width, (l) length between the anal and pelvic fins, and (m) pectoral fin width. Significance in violin plots is based on ANOVA analysis followed by Tukeys HSD (data in Table S2; p-values indicated by * <0.05, ** <0.01, *** <0.005). Genome-wide significance in QTL scans is indicated at the 5% (solid line) and 10% (dashed line) level. Details of QTL scans are in Table S1. QTL scans by chromosome are in Figure S4.

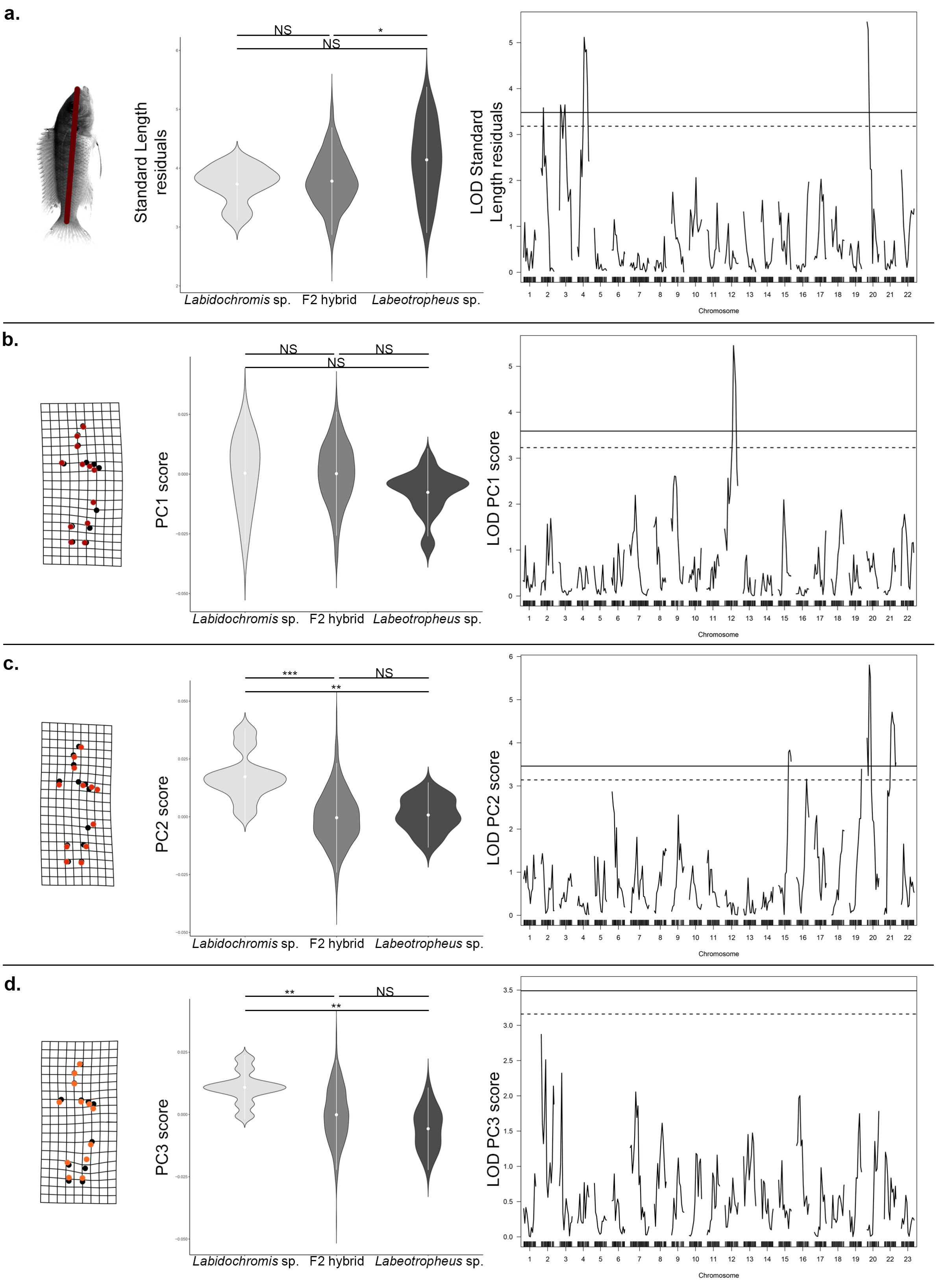
a.
NS
*
NS
Standard Length residuals
Labidochromis sp.
F2 hybrid
Labeotropheus sp.
LOD Standard Length residuals
Chromosome
b.
NS
NS
NS
PC1 score
Labidochromis sp.
F2 hybrid
Labeotropheus sp.
LOD PC1 score
Chromosome
c.
***
NS
**
PC2 score
Labidochromis sp.
F2 hybrid
Labeotropheus sp.
LOD PC2 score
Chromosome
d.
**
NS
**
PC3 score
Labidochromis sp.
F2 hybrid
Labeotropheus sp.
LOD PC3 score
Chromosome

**e.**

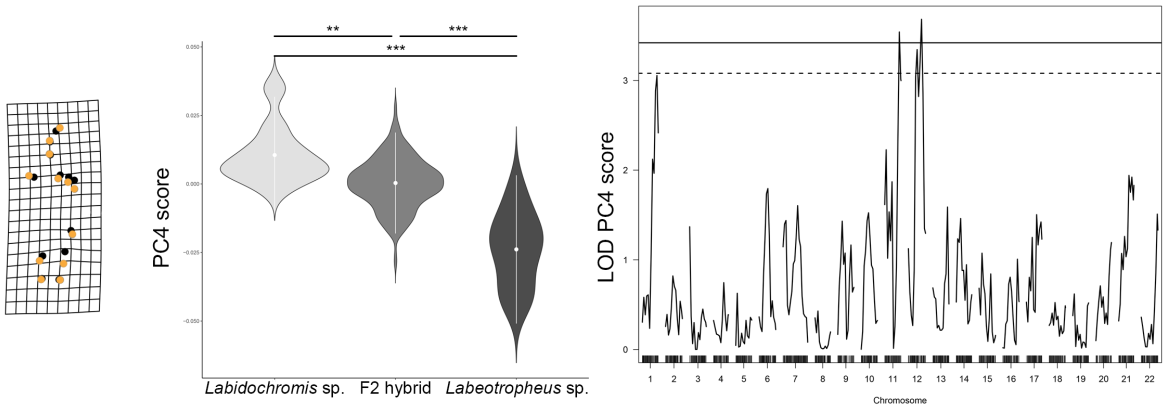
**
***
***
PC4 score
*Labidochromis* sp.
F2 hybrid
*Labeotropheus* sp.
LOD PC4 score
Chromosome

**f.**

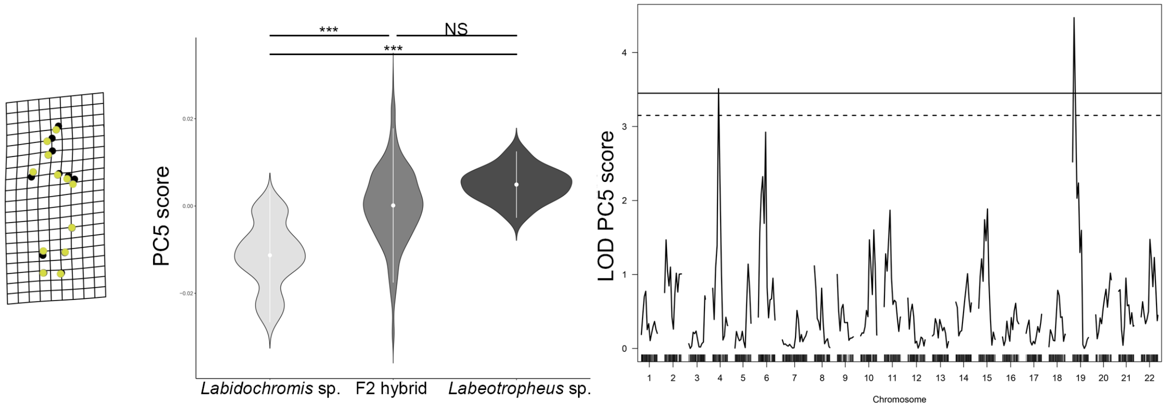
***
NS
***
PC5 score
*Labidochromis* sp.
F2 hybrid
*Labeotropheus* sp.
LOD PC5 score
Chromosome

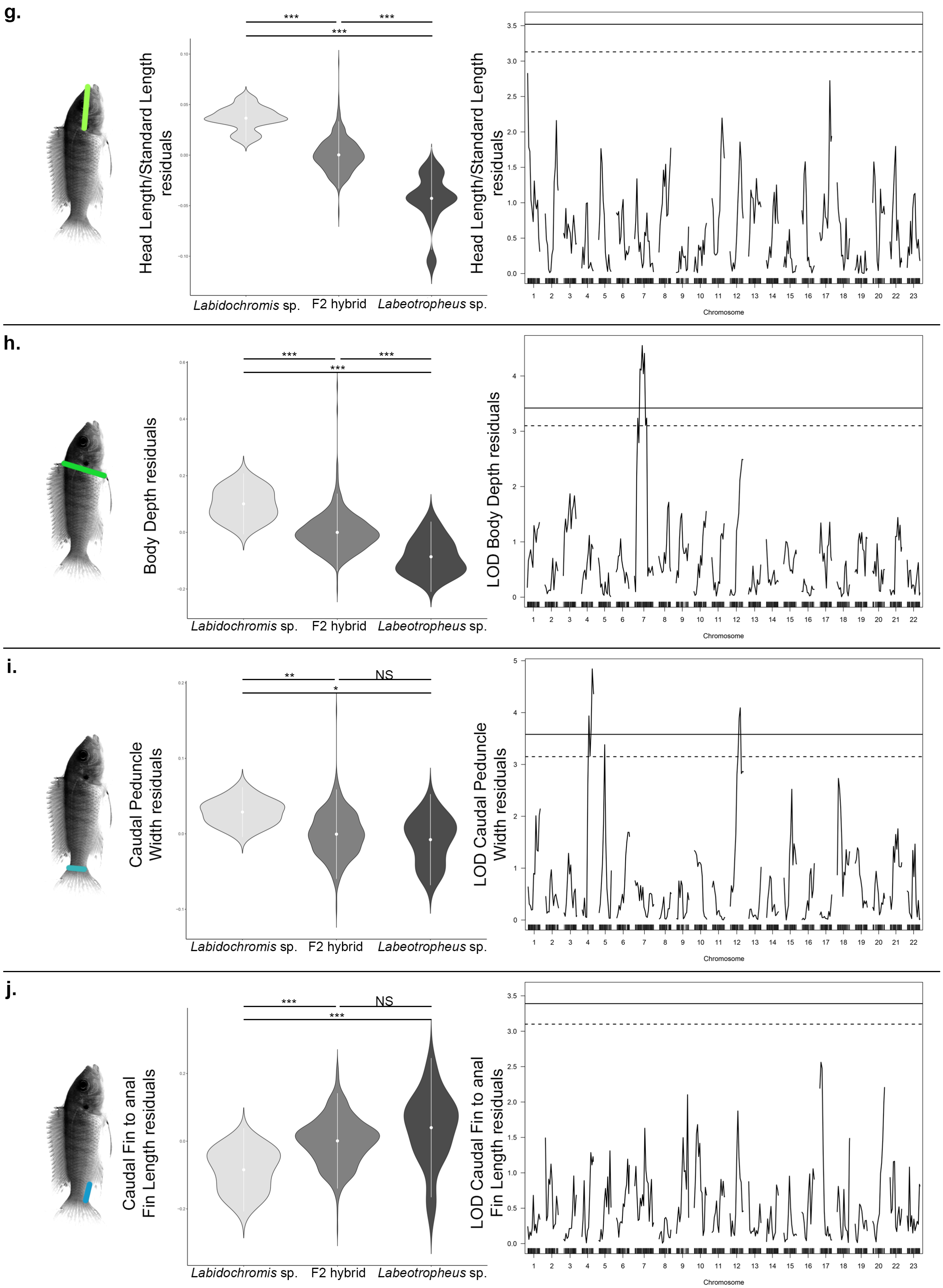
g.
h.
i.
j.
***
*
**
NS
Head Length/Standard Length residuals
LOD Head Length/Standard Length residuals
Body Depth residuals
LOD Body Depth residuals
Caudal Peduncle Width residuals
LOD Caudal Peduncle Width residuals
Caudal Fin to anal Fin Length residuals
LOD Caudal Fin to anal Fin Length residuals
Labidochromis sp.
F2 hybrid
Labeotropheus sp.
Chromosome

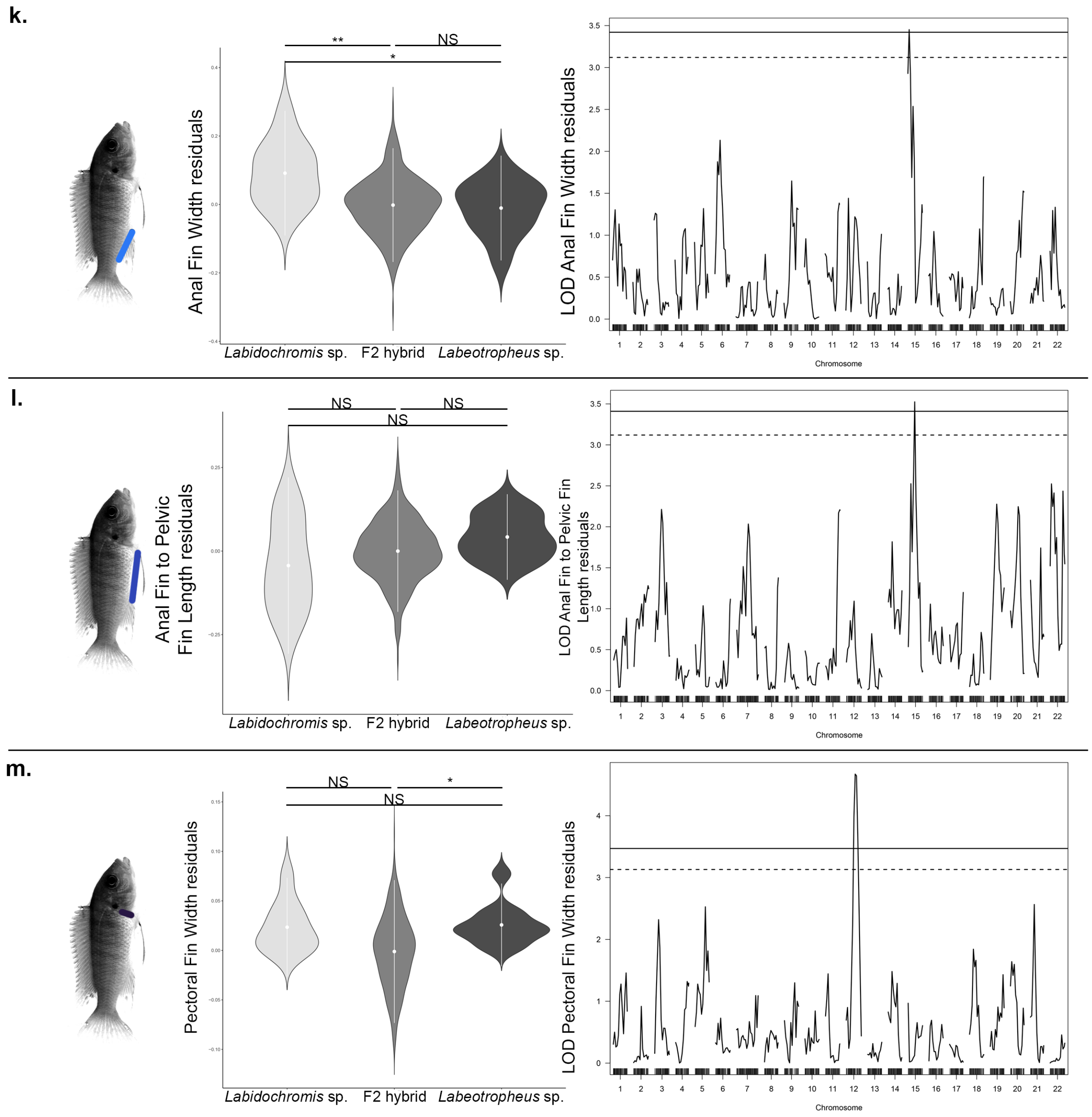


**Figure S2. Phenotypic measures and QTL scans within the *Labidochromis* x *Labeotropheus* cross.** Phenotypic measures are indicated by illustration with colors matching Figures 2 and S3 and include (a) standard length, (b) PC1 shape, (c) PC2 shape, (d) PC3 shape, (e) PC4 shape, (f) PC5 shape, (g) head proportion, measured as head length/standard length, (h) body depth, (i) caudal peduncle width, (j) length between the caudal and anal fins, (k) anal fin width, (l) length between the anal and pelvic fins, and (m) pectoral fin width. Significance in violin plots is based on ANOVA analysis followed by Tukeys HSD (data in Table S2; p-values indicated by * <0.05, ** <0.01, *** <0.005). Genome-wide significance in QTL scans is indicated at the 5% (solid line) and 10% (dashed line) level. Details of QTL scans are in Table S1. QTL scans by chromosome are in Figure S5.

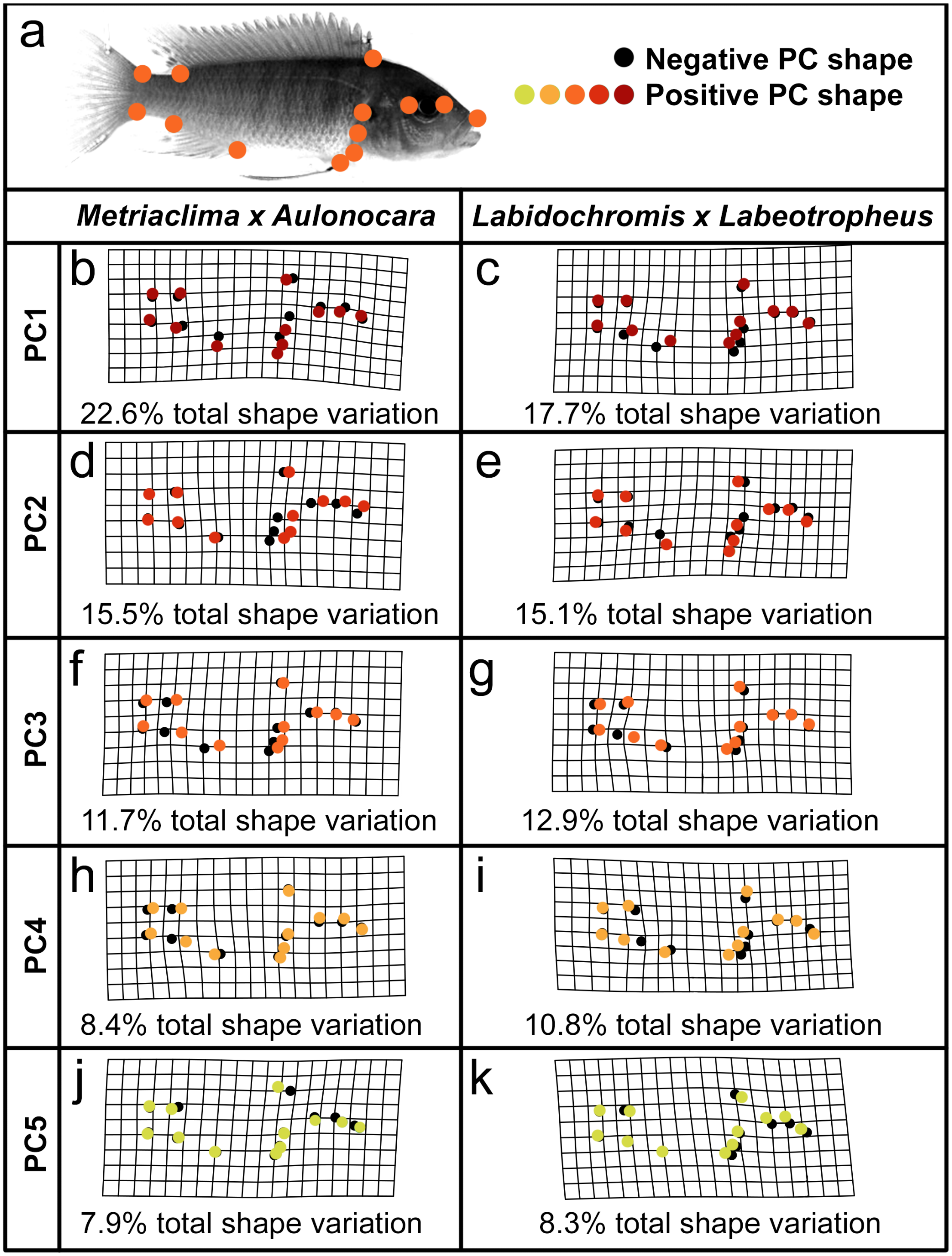


**Figure S3. Shapes indicated by PC scores following geometric morphometric analysis.** Landmarks used in analysis are as indicated in (a) and described in Figure 2. Shape illustrated by negative PC scores are in black dots, while colored dots indicate shape described by positive PC scores for (b-c) PC1 scores, (d-e) PC2 scores, (f-g) PC3 scores, (h-i) PC4 scores, and (j-k) PC5 scores. Total phenotypic variation explained by each PC in both crosses is indicated.

a.

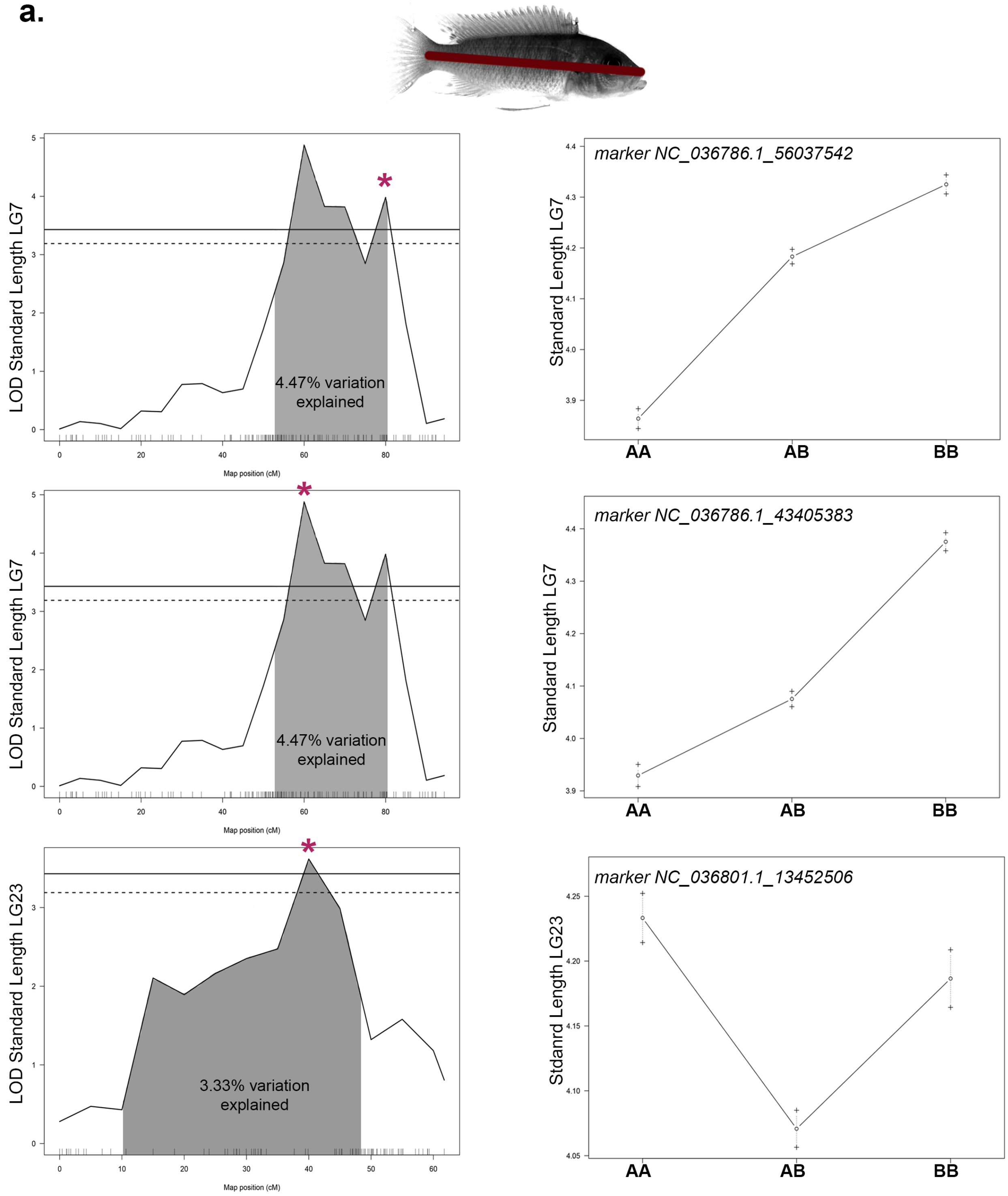
LOD Standard Length LG7
4.47% variation explained
Map position (cM)
marker NC_036786.1_56037542
Standard Length LG7
AA
AB
BB
LOD Standard Length LG7
4.47% variation explained
Map position (cM)
marker NC_036786.1_43405383
Standard Length LG7
AA
AB
BB
LOD Standard Length LG23
3.33% variation explained
Map position (cM)
marker NC_036801.1_13452506
Stdanrd Length LG23
AA
AB
BB

**b.**

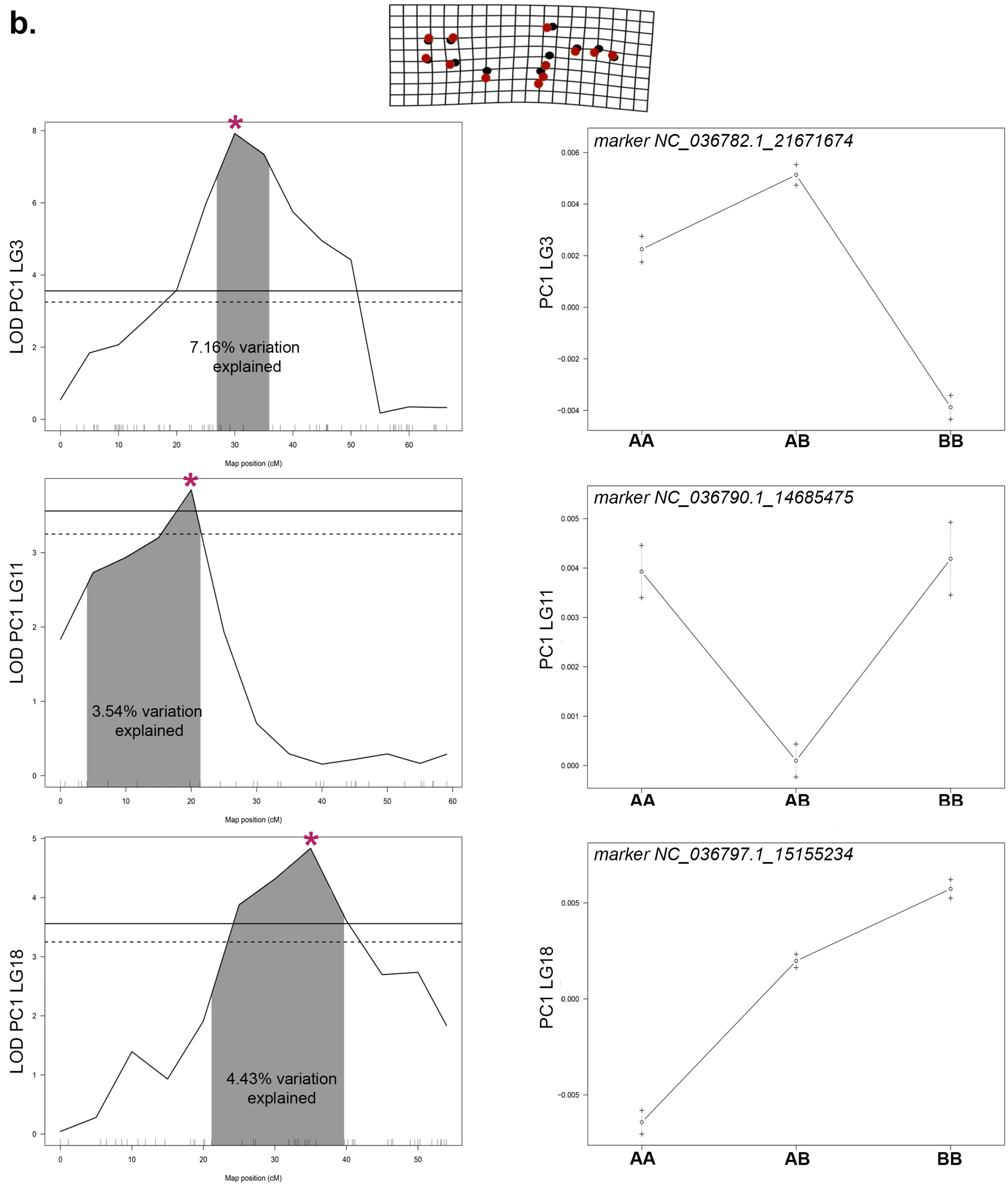

LOD PC1 LG3
7.16% variation explained
Map position (cM)
marker NC_036782.1_21671674
PC1 LG3
AA
AB
BB
LOD PC1 LG11
3.54% variation explained
Map position (cM)
marker NC_036790.1_14685475
PC1 LG11
AA
AB
BB
LOD PC1 LG18
4.43% variation explained
Map position (cM)
marker NC_036797.1_15155234
PC1 LG18
AA
AB
BB

c.

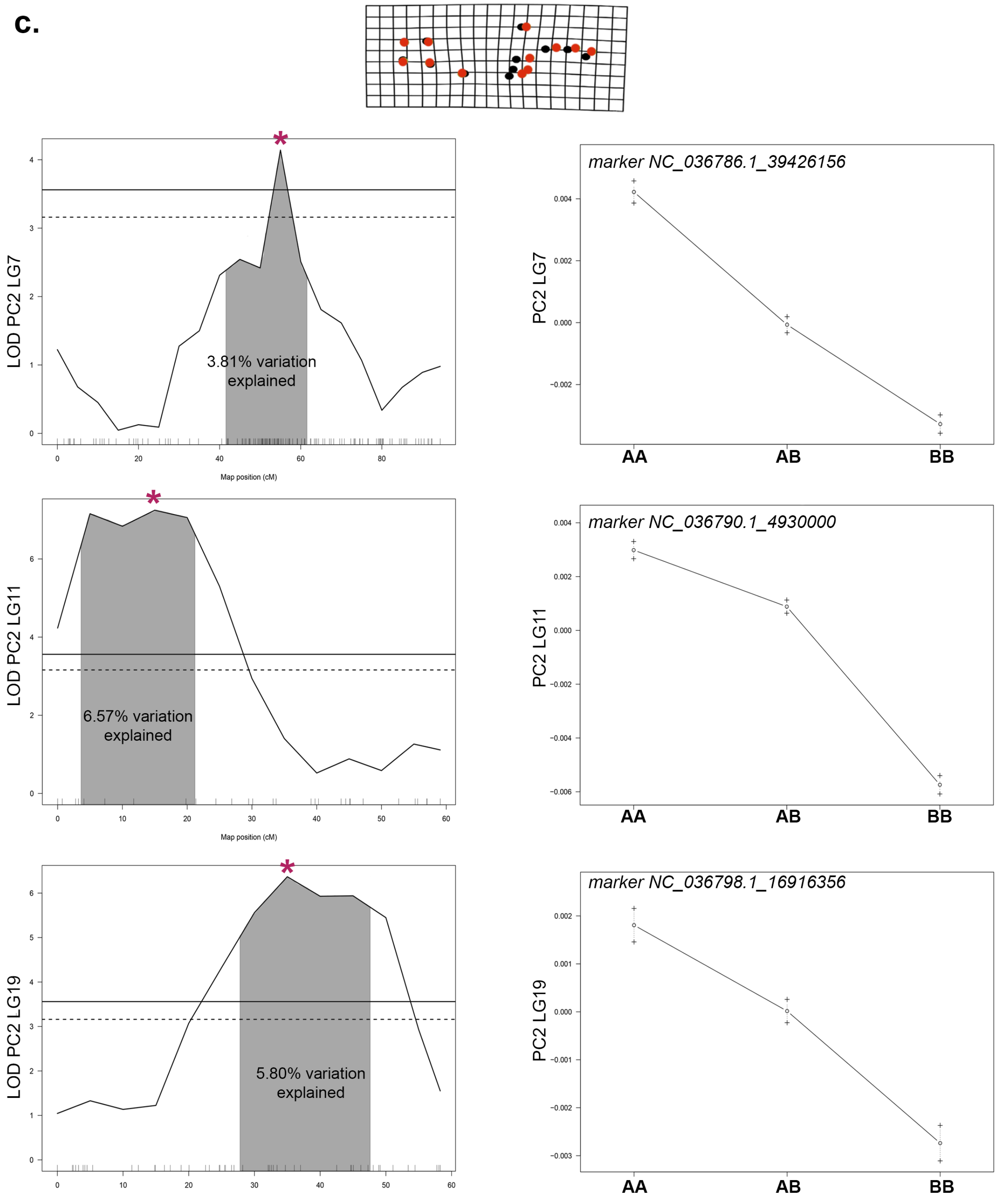
LOD PC2 LG7
3.81% variation explained
Map position (cM)
marker NC_036786.1_39426156
PC2 LG7
AA
AB
BB
LOD PC2 LG11
6.57% variation explained
Map position (cM)
marker NC_036790.1_4930000
PC2 LG11
AA
AB
BB
LOD PC2 LG19
5.80% variation explained
marker NC_036798.1_16916356
PC2 LG19
AA
AB
BB

**d.**

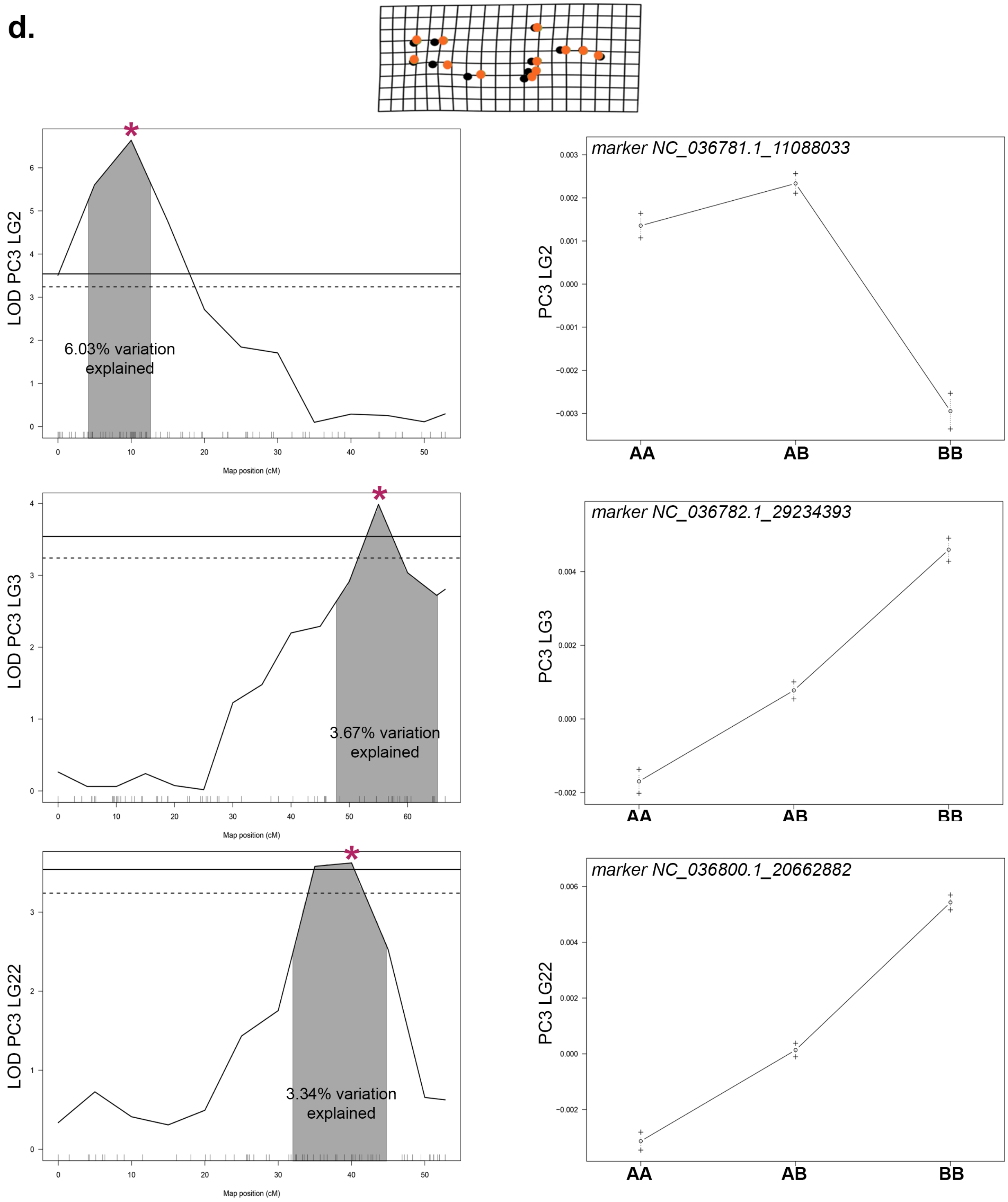

*
LOD PC3 LG2
6.03% variation explained
Map position (cM)
marker NC_036781.1_11088033
PC3 LG2
AA
AB
BB
*
LOD PC3 LG3
3.67% variation explained
Map position (cM)
marker NC_036782.1_29234393
PC3 LG3
AA
AB
BB
*
LOD PC3 LG22
3.34% variation explained
Map position (cM)
marker NC_036800.1_20662882
PC3 LG22
AA
AB
BB

e.

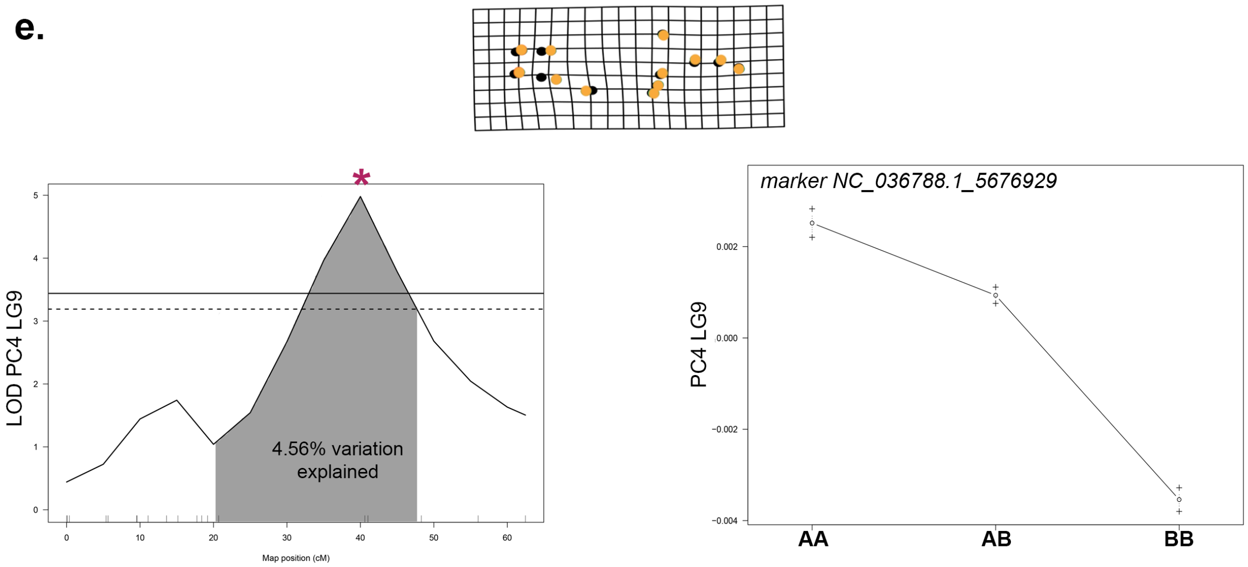
LOD PC4 LG9
4.56% variation explained
Map position (cM)
marker NC_036788.1_5676929
PC4 LG9
AA
AB
BB

f.

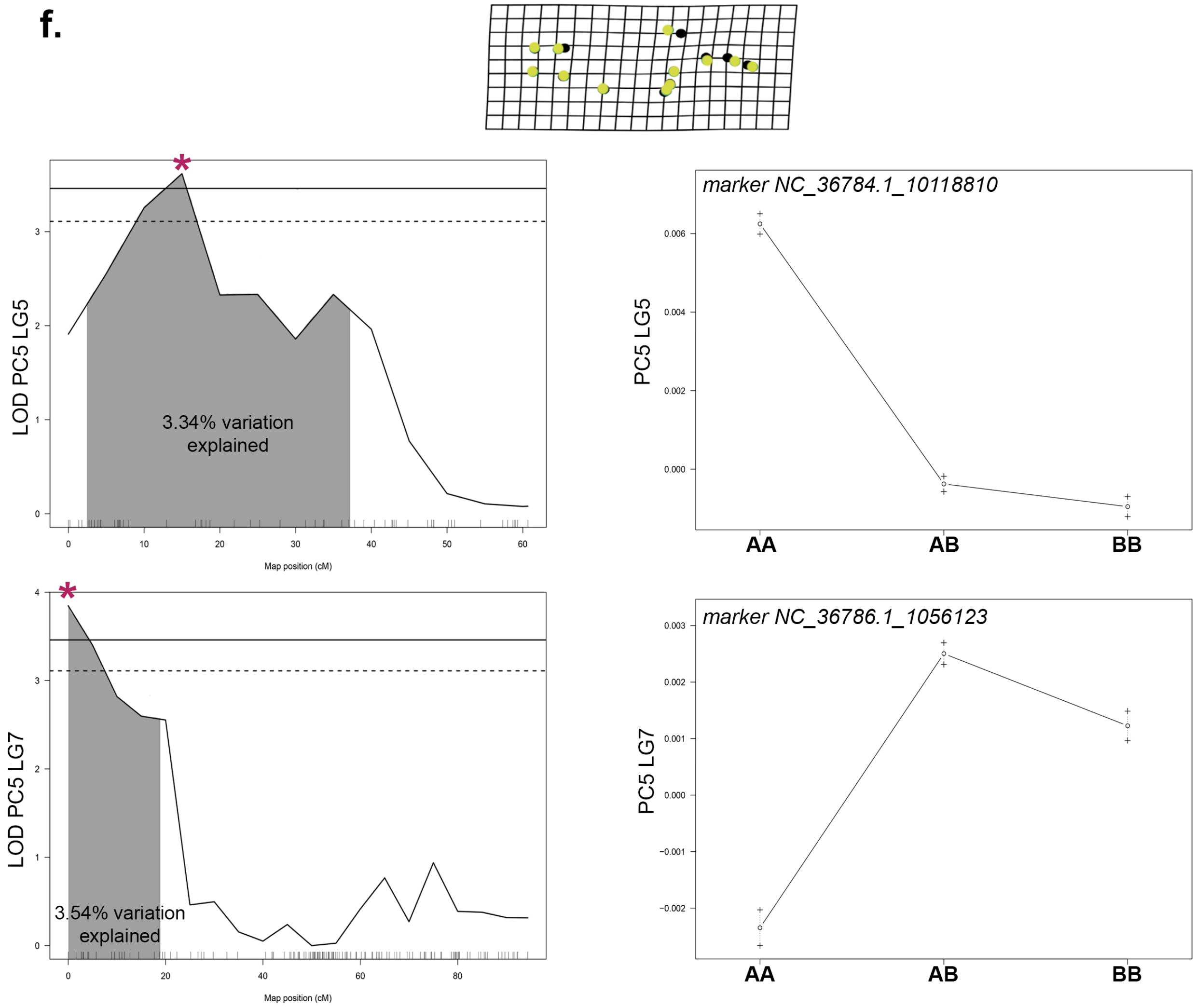
LOD PC5 LG5
3.34% variation explained
Map position (cM)
marker NC_36784.1_10118810
PC5 LG5
AA
AB
BB
LOD PC5 LG7
3.54% variation explained
Map position (cM)
marker NC_36786.1_1056123
PC5 LG7
AA
AB
BB

**f.**

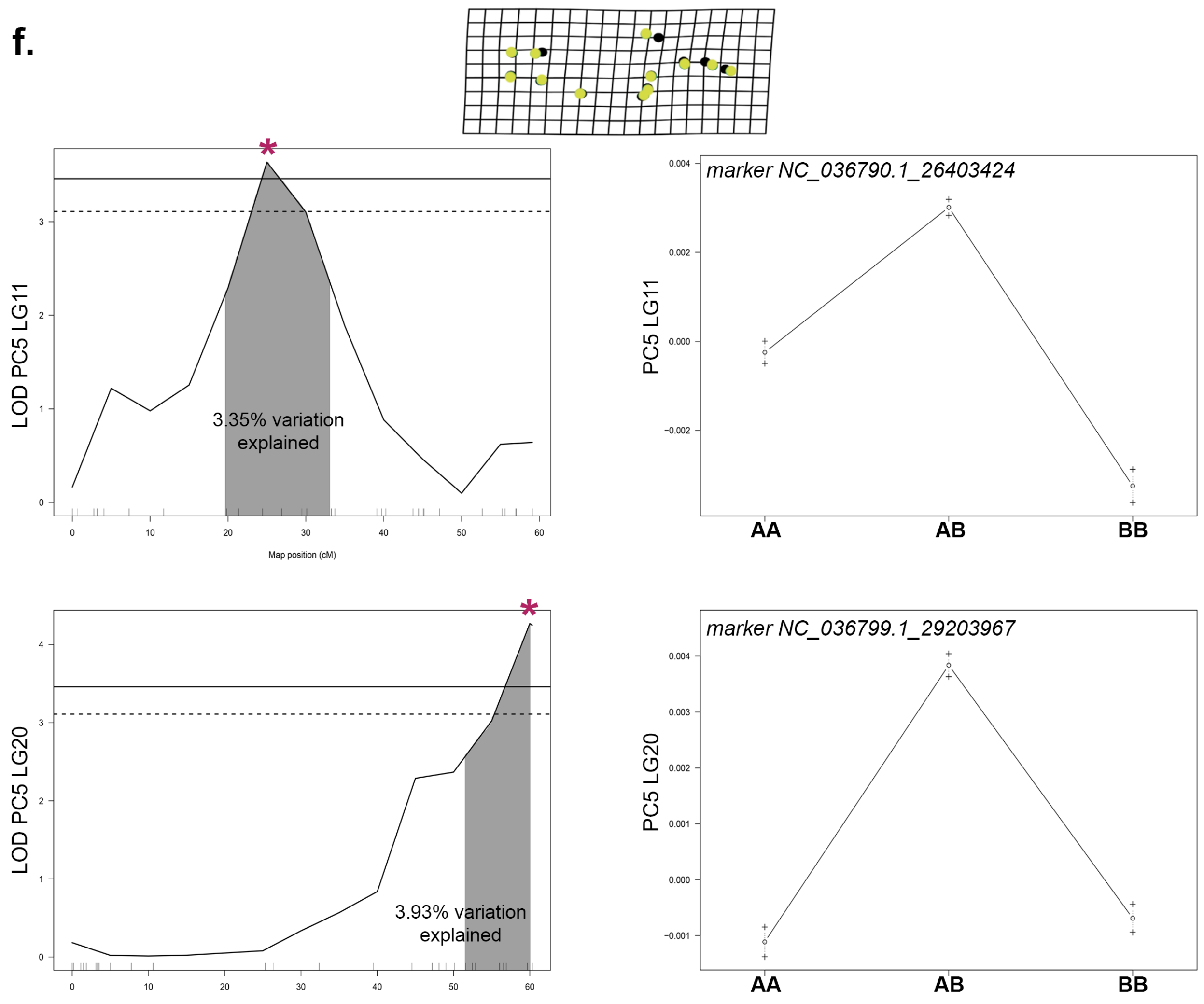

LOD PC5 LG11
3.35% variation explained
Map position (cM)
marker NC_036790.1_26403424
PC5 LG11
AA
AB
BB
LOD PC5 LG20
3.93% variation explained
marker NC_036799.1_29203967
PC5 LG20
AA
AB
BB

g.

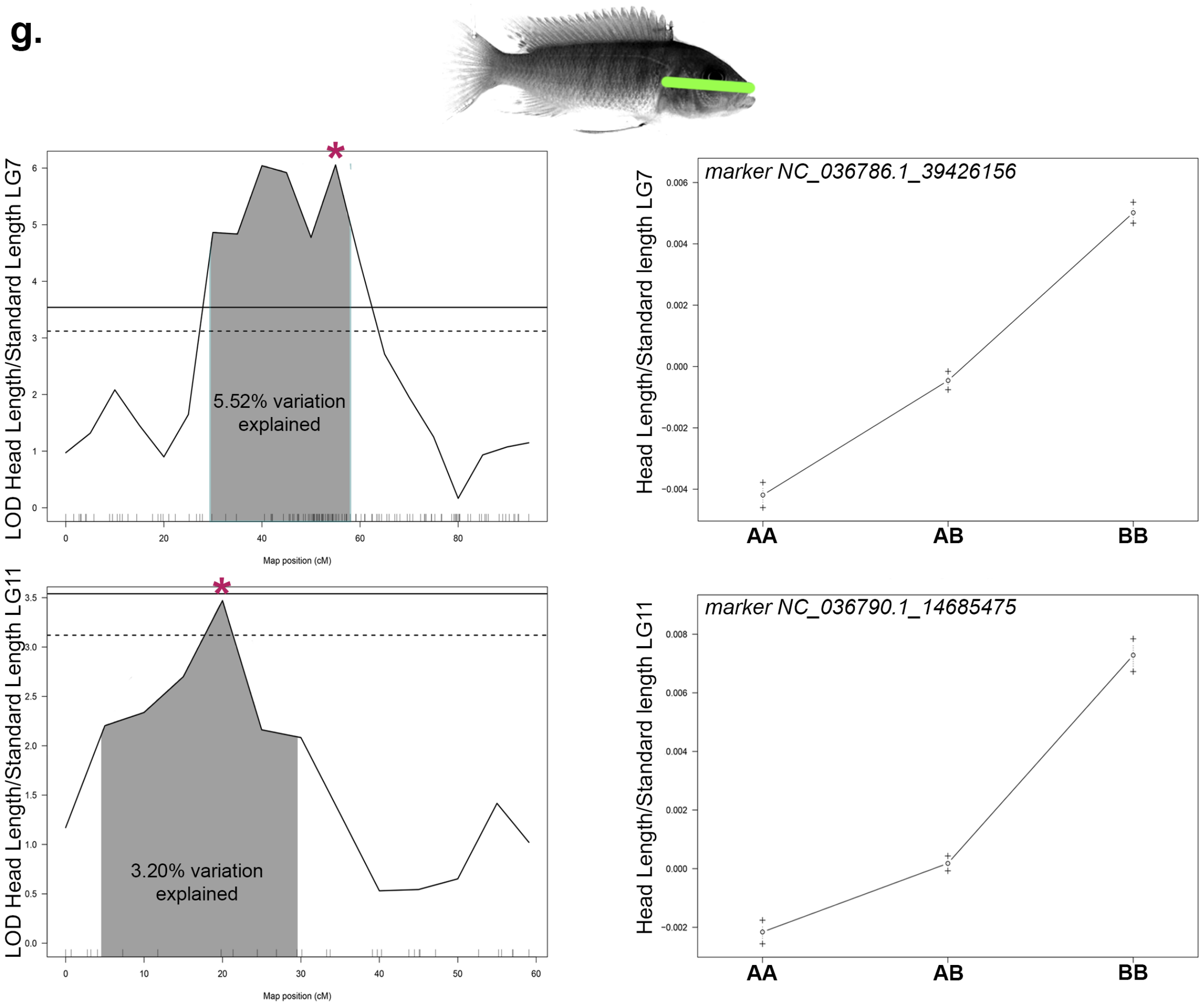
LOD Head Length/Standard Length LG7
5.52% variation explained
Map position (cM)
marker NC_036786.1_39426156
Head Length/Standard length LG7
AA
AB
BB
LOD Head Length/Standard Length LG11
3.20% variation explained
Map position (cM)
marker NC_036790.1_14685475
Head Length/Standard length LG11
AA
AB
BB

**h.**

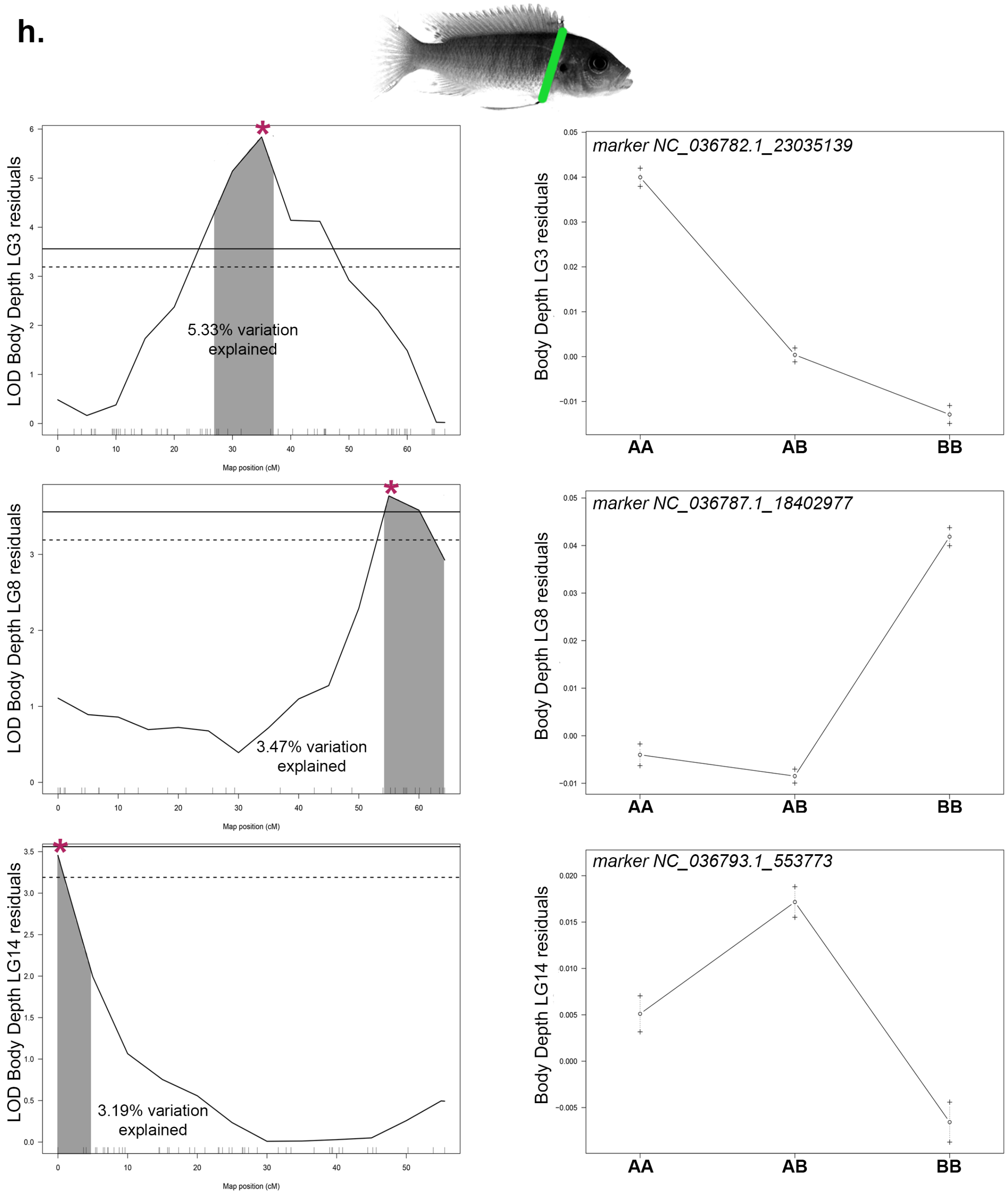
LOD Body Depth LG3 residuals
5.33% variation explained
Map position (cM)
marker NC_036782.1_23035139
Body Depth LG3 residuals
AA
AB
BB
LOD Body Depth LG8 residuals
3.47% variation explained
Map position (cM)
marker NC_036787.1_18402977
Body Depth LG8 residuals
AA
AB
BB
LOD Body Depth LG14 residuals
3.19% variation explained
Map position (cM)
marker NC_036793.1_553773
Body Depth LG14 residuals
AA
AB
BB

**h.**

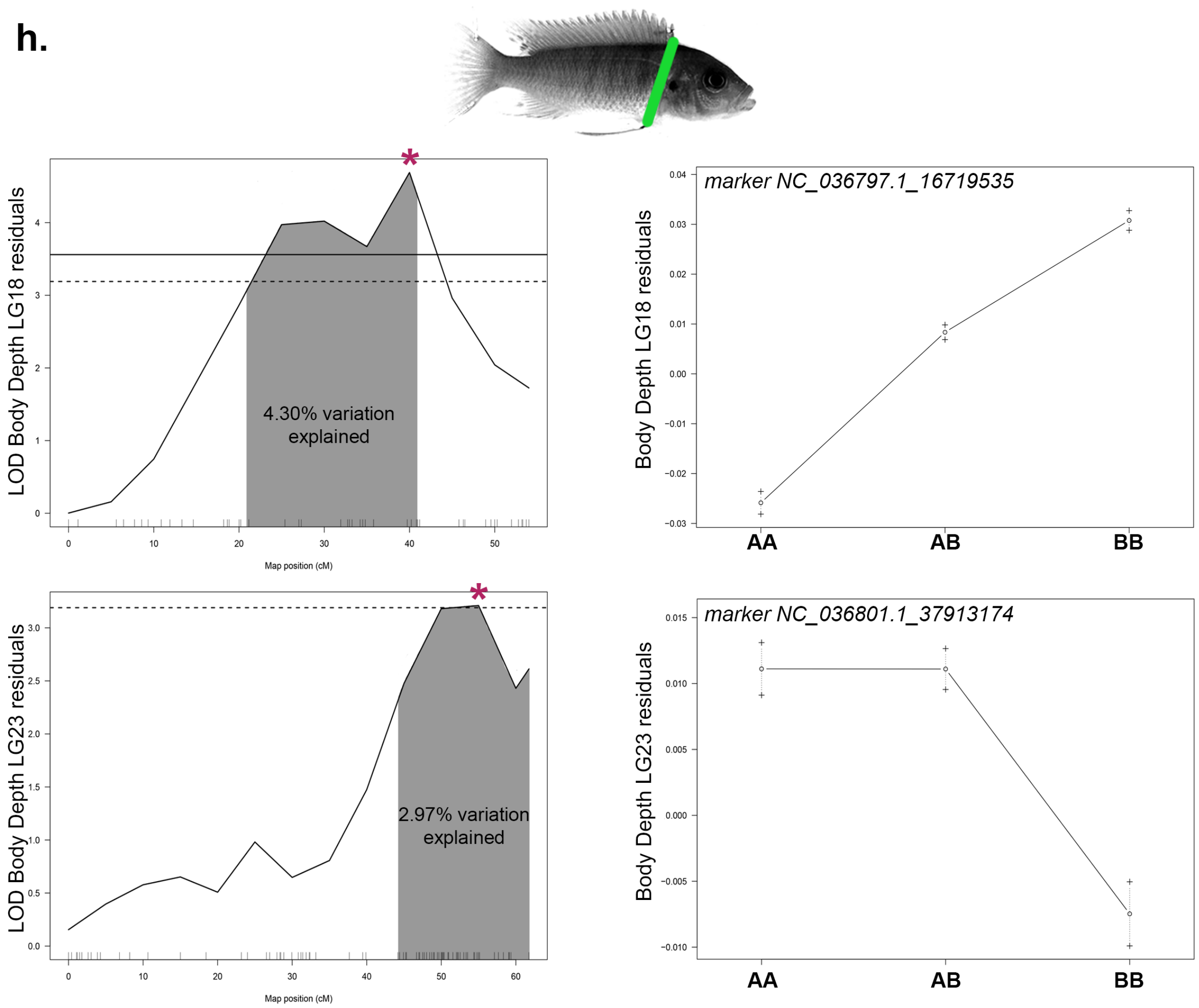

LOD Body Depth LG18 residuals
4.30% variation explained
Map position (cM)
marker NC_036797.1_16719535
Body Depth LG18 residuals
AA
AB
BB
LOD Body Depth LG23 residuals
2.97% variation explained
Map position (cM)
marker NC_036801.1_37913174
Body Depth LG23 residuals
AA
AB
BB

**i.**

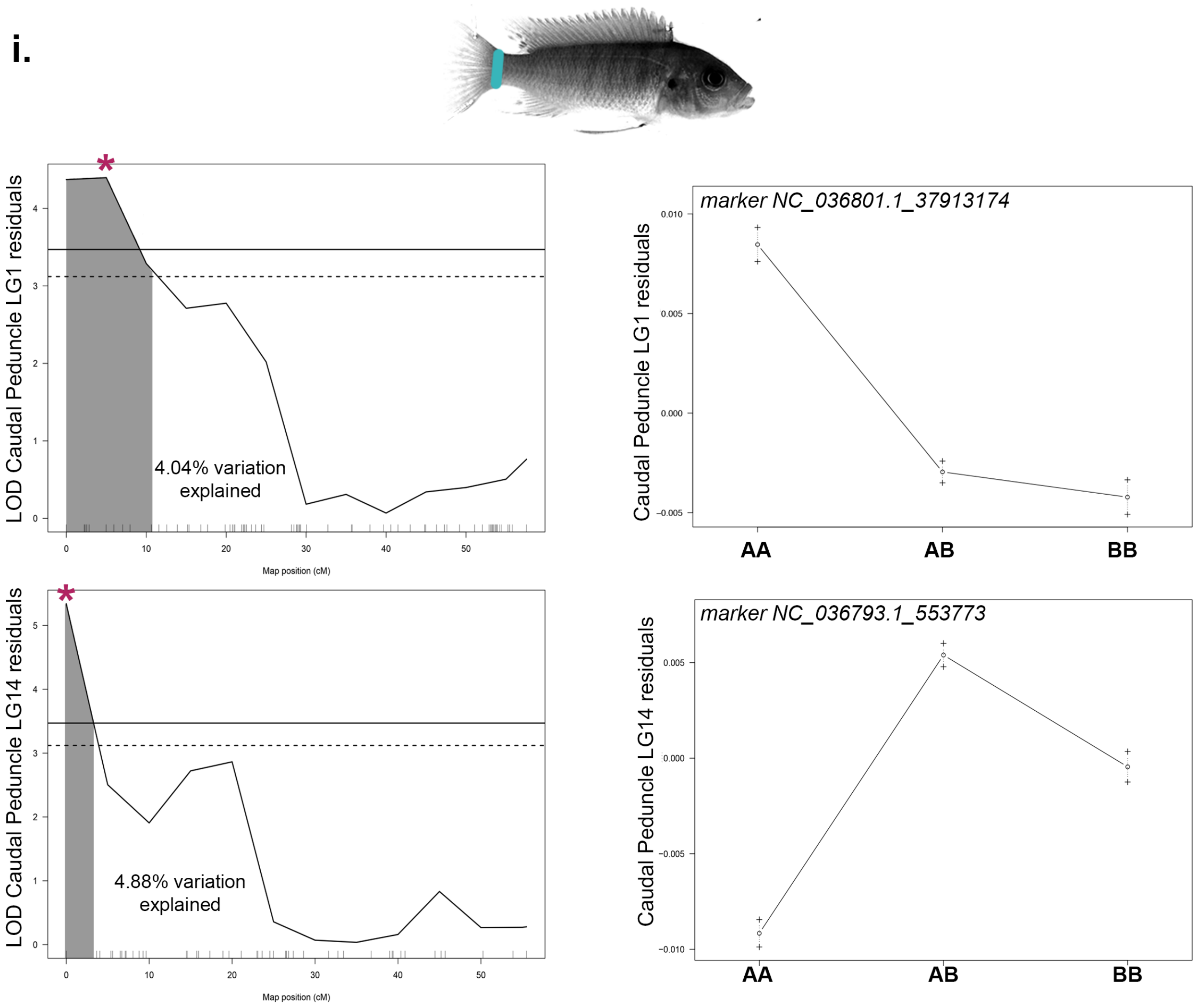

LOD Caudal Peduncle LG1 residuals
4.04% variation explained
Map position (cM)
marker NC_036801.1_37913174
Caudal Peduncle LG1 residuals
AA
AB
BB
LOD Caudal Peduncle LG14 residuals
4.88% variation explained
Map position (cM)
marker NC_036793.1_553773
Caudal Peduncle LG14 residuals
AA
AB
BB

**j.**

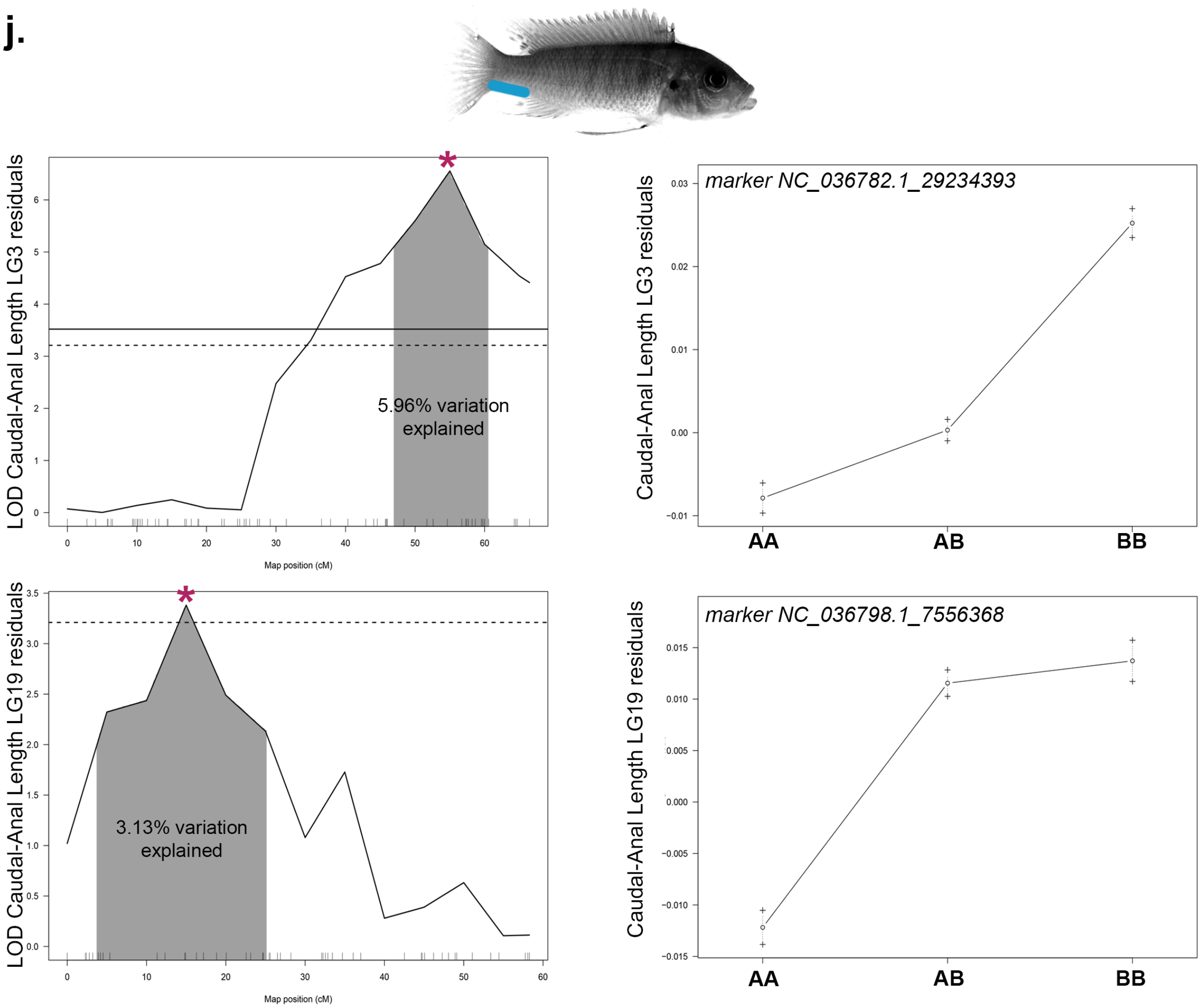

5.96% variation explained
LOD Caudal-Anal Length LG3 residuals
Map position (cM)
marker NC_036782.1_29234393
Caudal-Anal Length LG3 residuals
AA
AB
BB
3.13% variation explained
LOD Caudal-Anal Length LG19 residuals
Map position (cM)
marker NC_036798.1_7556368
Caudal-Anal Length LG19 residuals
AA
AB
BB

**k.**

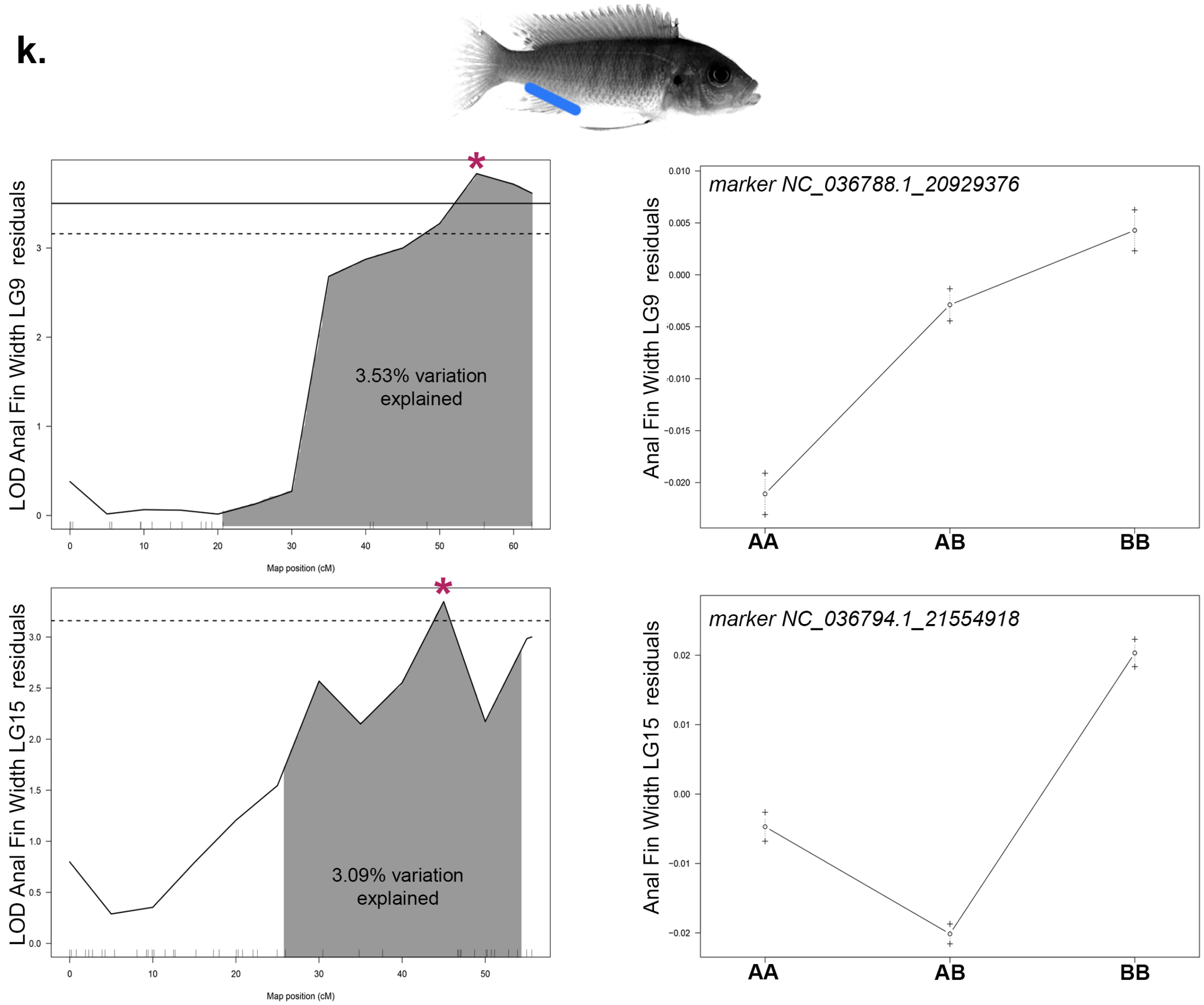

LOD Anal Fin Width LG9 residuals
3.53% variation explained
Map position (cM)
marker NC_036788.1_20929376
Anal Fin Width LG9 residuals
AA
AB
BB
LOD Anal Fin Width LG15 residuals
3.09% variation explained
Map position (cM)
marker NC_036794.1_21554918
Anal Fin Width LG15 residuals
AA
AB
BB


**l.**

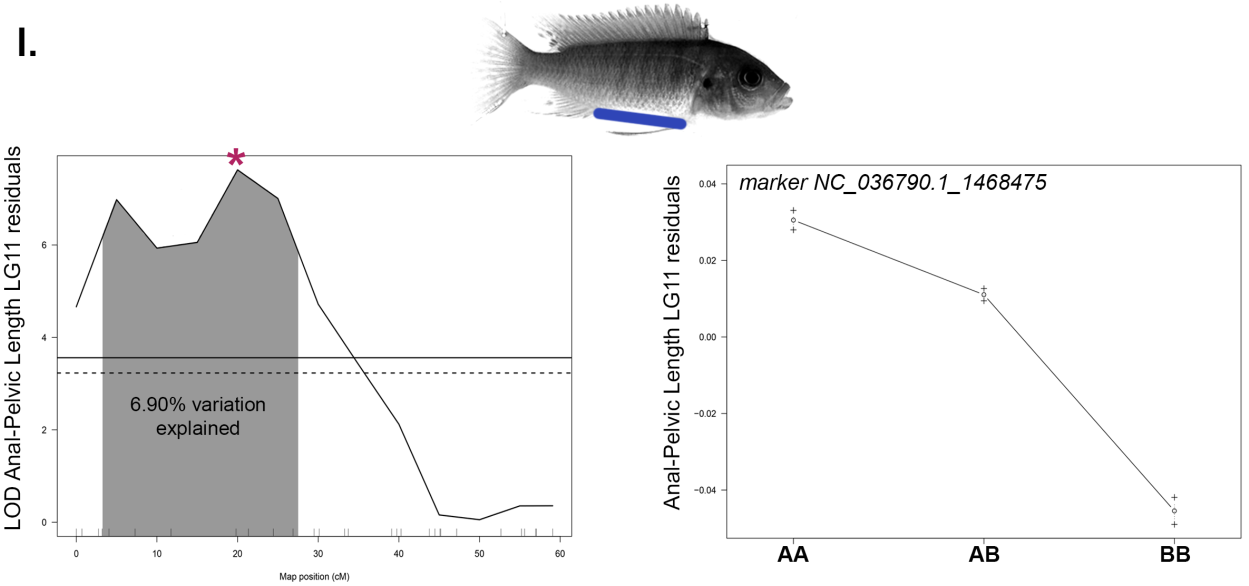

LOD Anal-Pelvic Length LG11 residuals
6.90% variation explained
Map position (cM)
marker NC_036790.1_1468475
Anal-Pelvic Length LG11 residuals
AA
AB
BB

m.

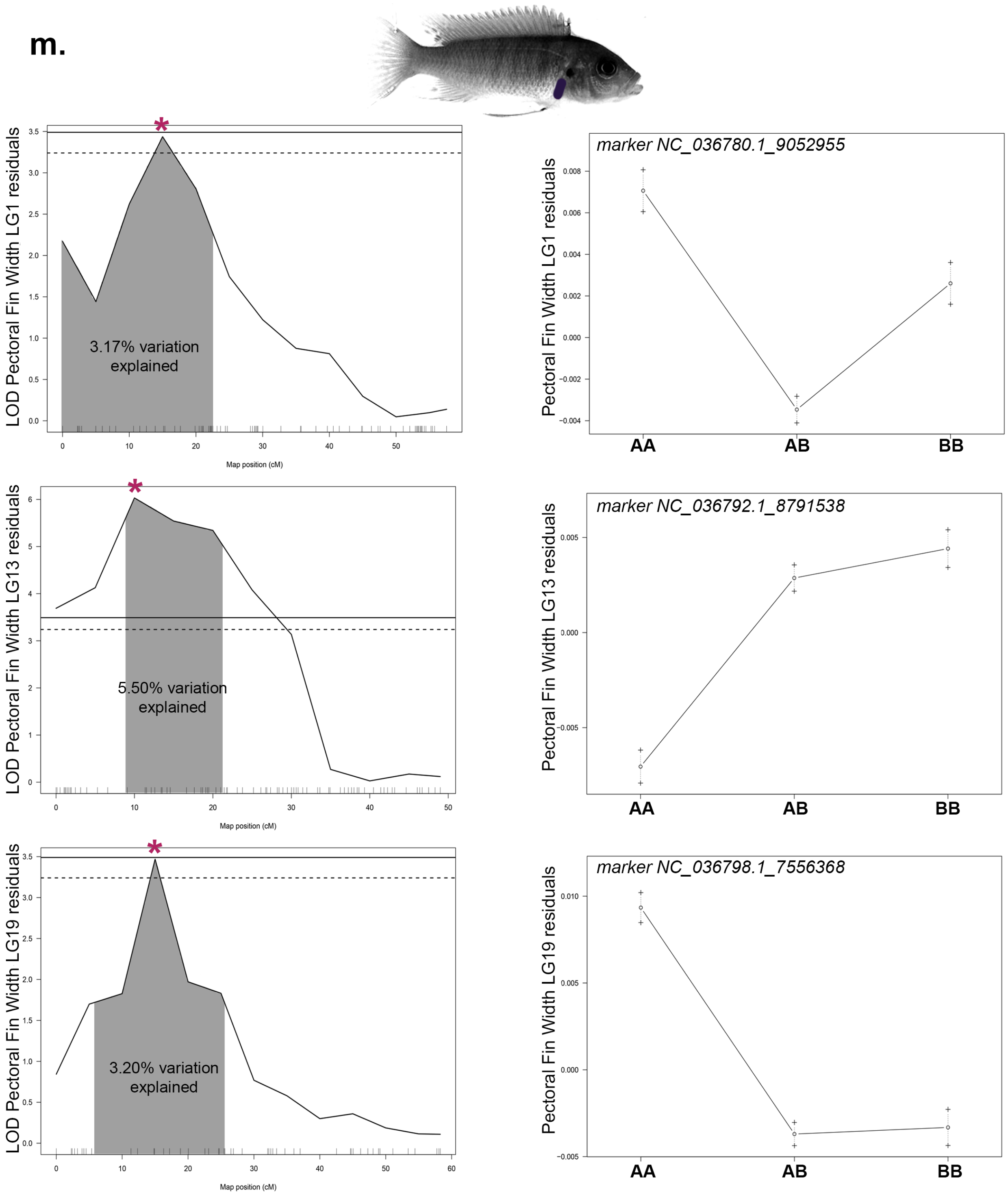


**Figure S4. QTL scans and allelic effects for significant QTL in *Metriaclima* x *Aulonocara* cross.** Phenotypic measures are indicated by illustration with colors matching Figures 2 and S3 and include (a) standard length, (b) PC1 shape, (c) PC2 shape, (d) PC3 shape, (e) PC4 shape, (f) PC5 shape, (g) head proportion, measured as head length/standard length, (h) body depth, (i)

caudal peduncle width, (j) length between the caudal and anal fins, (k) anal fin width, (l) length between the anal and pelvic fins, and (m) pectoral fin width. 95% confidence interval for QTL is indicated by shading, percent of total phenotypic variation explained by QTL is reported, and genome-wide significance is shown at the 5% (solid line) and 10% (dashed line) level. Details of QTL scan are in Table S1 and genome-wide visuals are in Figure S1. Allelic effects are shown for the position indicated by *. The A allele was inherited from the *Metriaclima* granddam and the B allele from the *Aulonocara* grandsire.

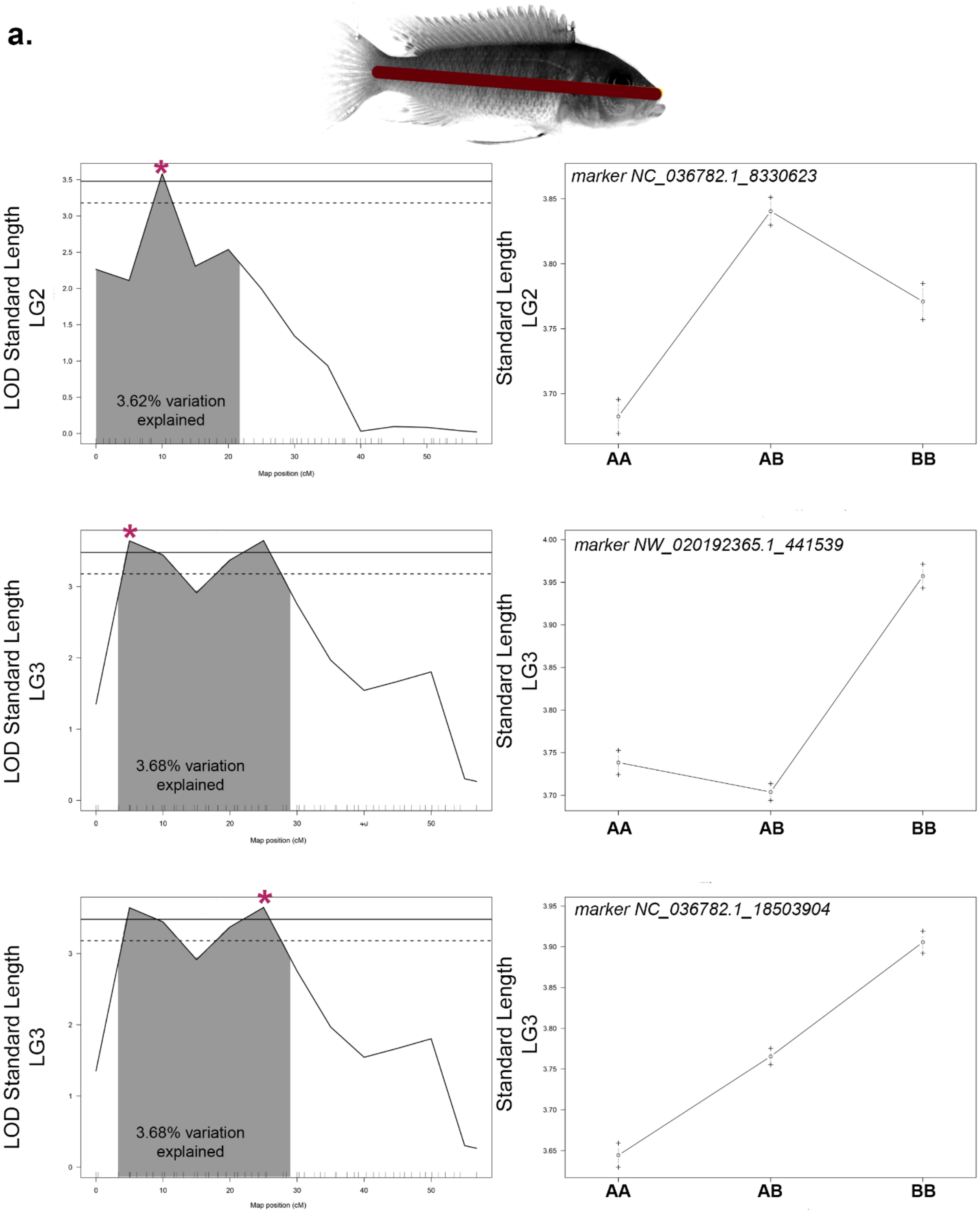
a.
LOD Standard Length LG2
3.62% variation explained
Map position (cM)
Standard Length LG2
marker NC_036782.1_8330623
AA
AB
BB
LOD Standard Length LG3
3.68% variation explained
Map position (cM)
Standard Length LG3
marker NW_020192365.1_441539
AA
AB
BB
LOD Standard Length LG3
3.68% variation explained
Map position (cM)
Standard Length LG3
marker NC_036782.1_18503904
AA
AB
BB

a.

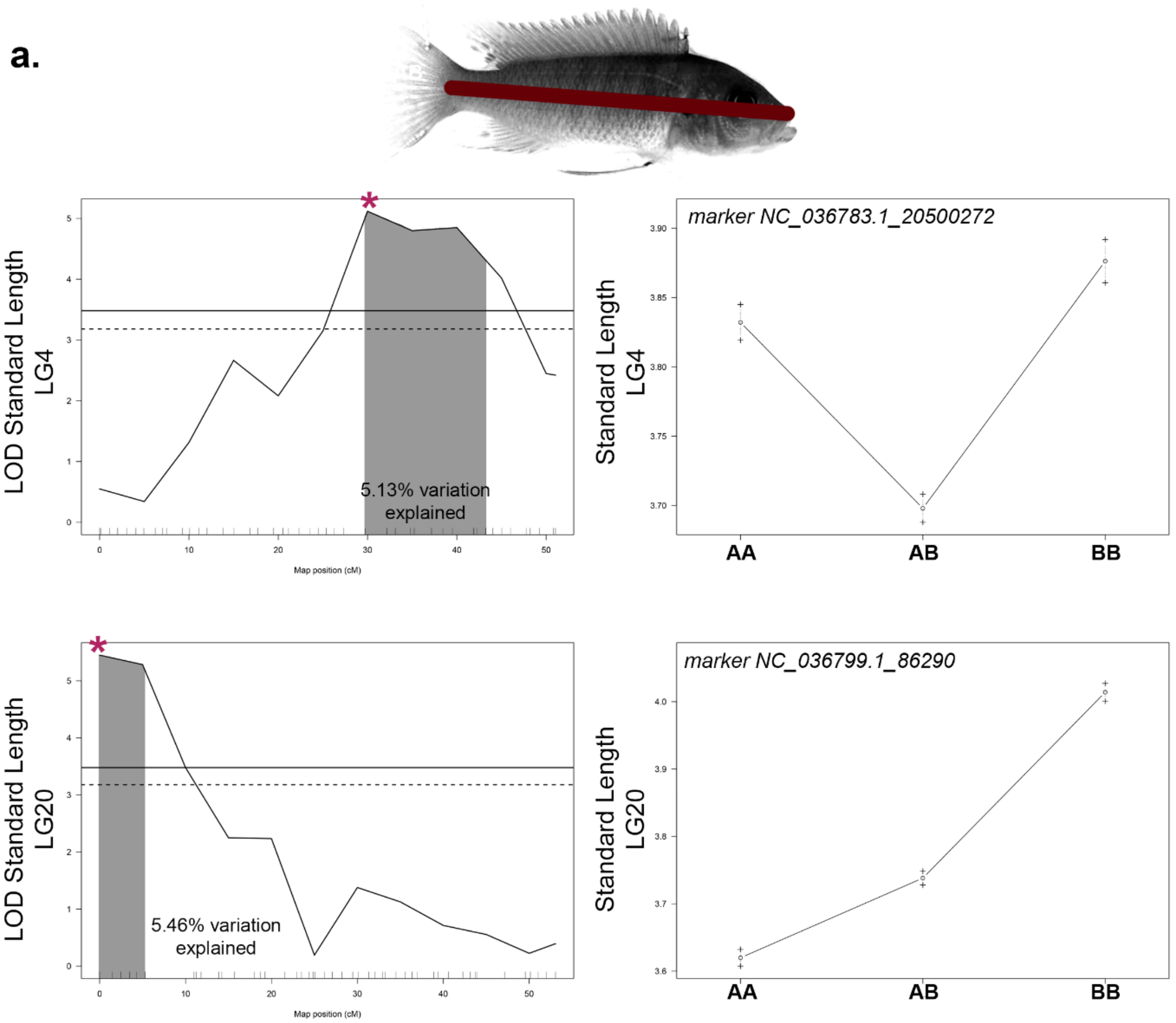

marker NC_036783.1_20500272
LOD Standard Length LG4
5.13% variation explained
Map position (cM)
Standard Length LG4
AA
AB
BB
marker NC_036799.1_86290
LOD Standard Length LG20
5.46% variation explained
Map position (cM)
Standard Length LG20
AA
AB
BB

**b.**

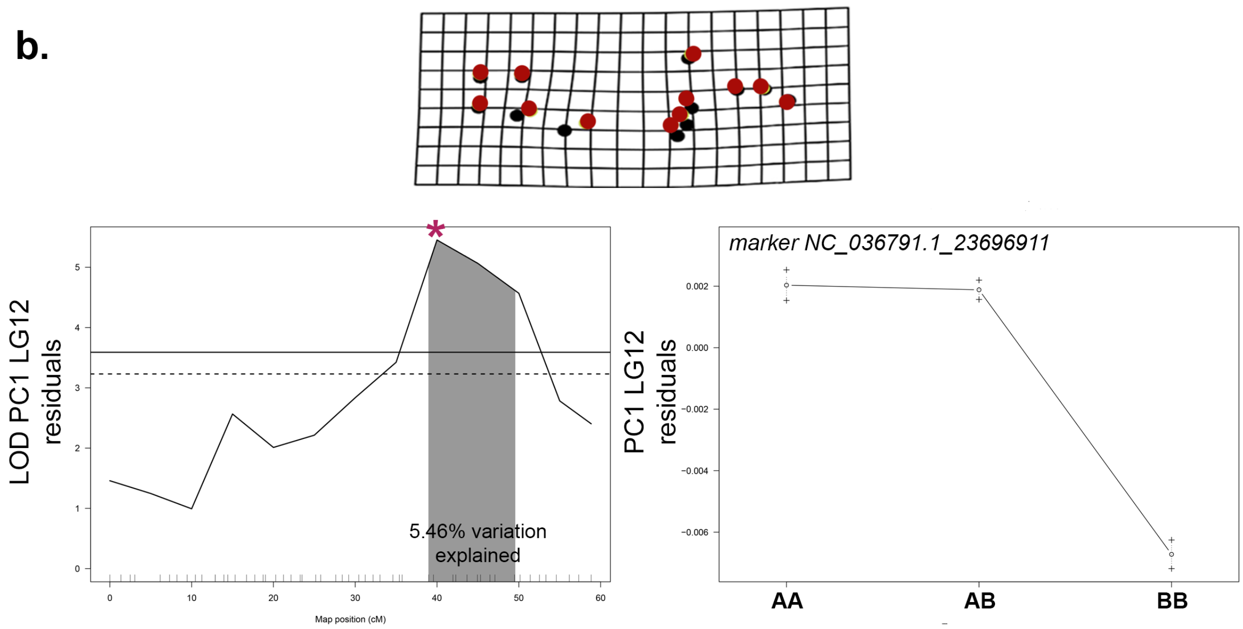

marker NC_036791.1_23696911
LOD PC1 LG12 residuals
5.46% variation explained
Map position (cM)
PC1 LG12 residuals
AA
AB
BB

**c.**

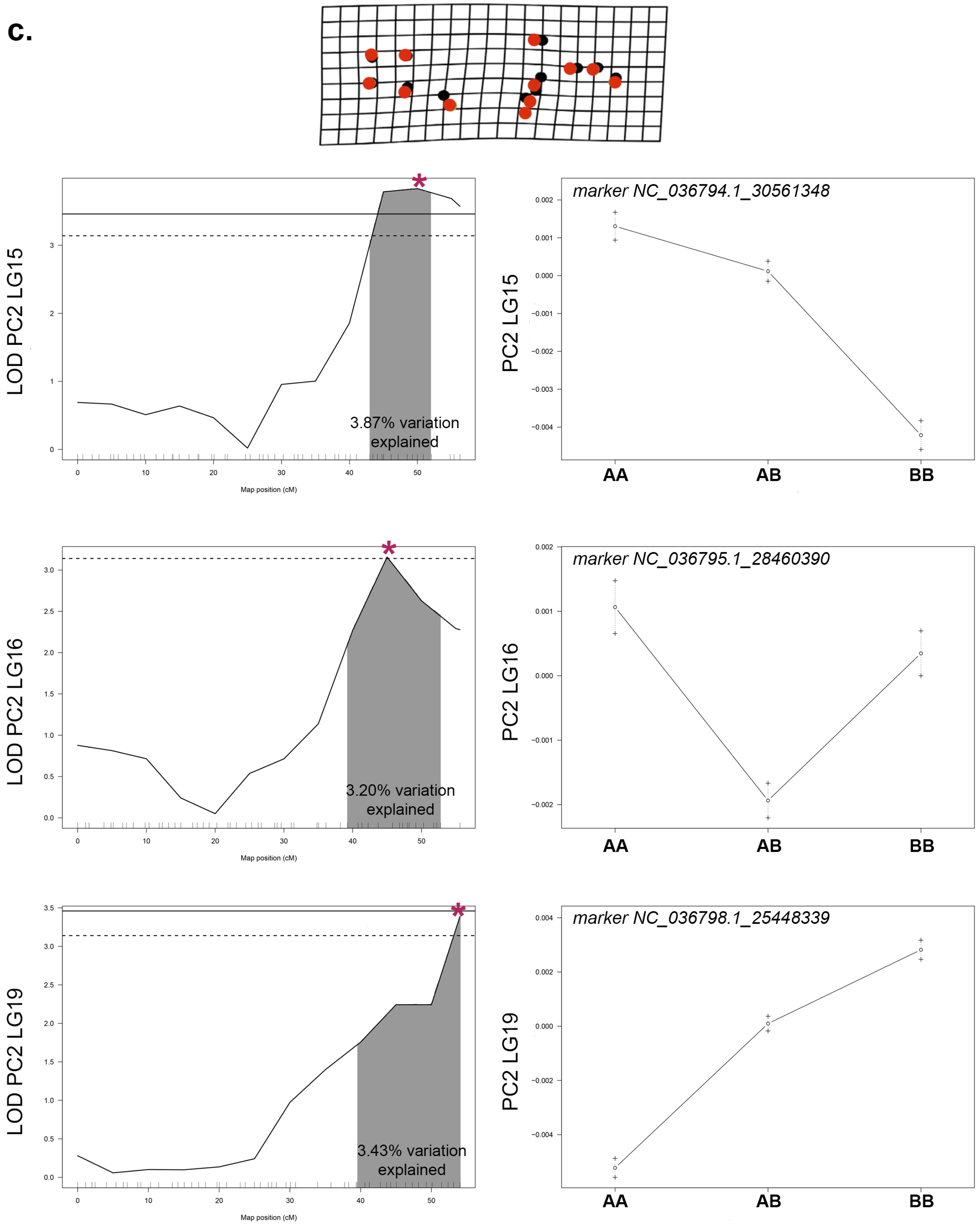

LOD PC2 LG15
3.87% variation explained
Map position (cM)
marker NC_036794.1_30561348
PC2 LG15
AA
AB
BB
LOD PC2 LG16
3.20% variation explained
Map position (cM)
marker NC_036795.1_28460390
PC2 LG16
AA
AB
BB
LOD PC2 LG19
3.43% variation explained
Map position (cM)
marker NC_036798.1_25448339
PC2 LG19
AA
AB
BB

c.

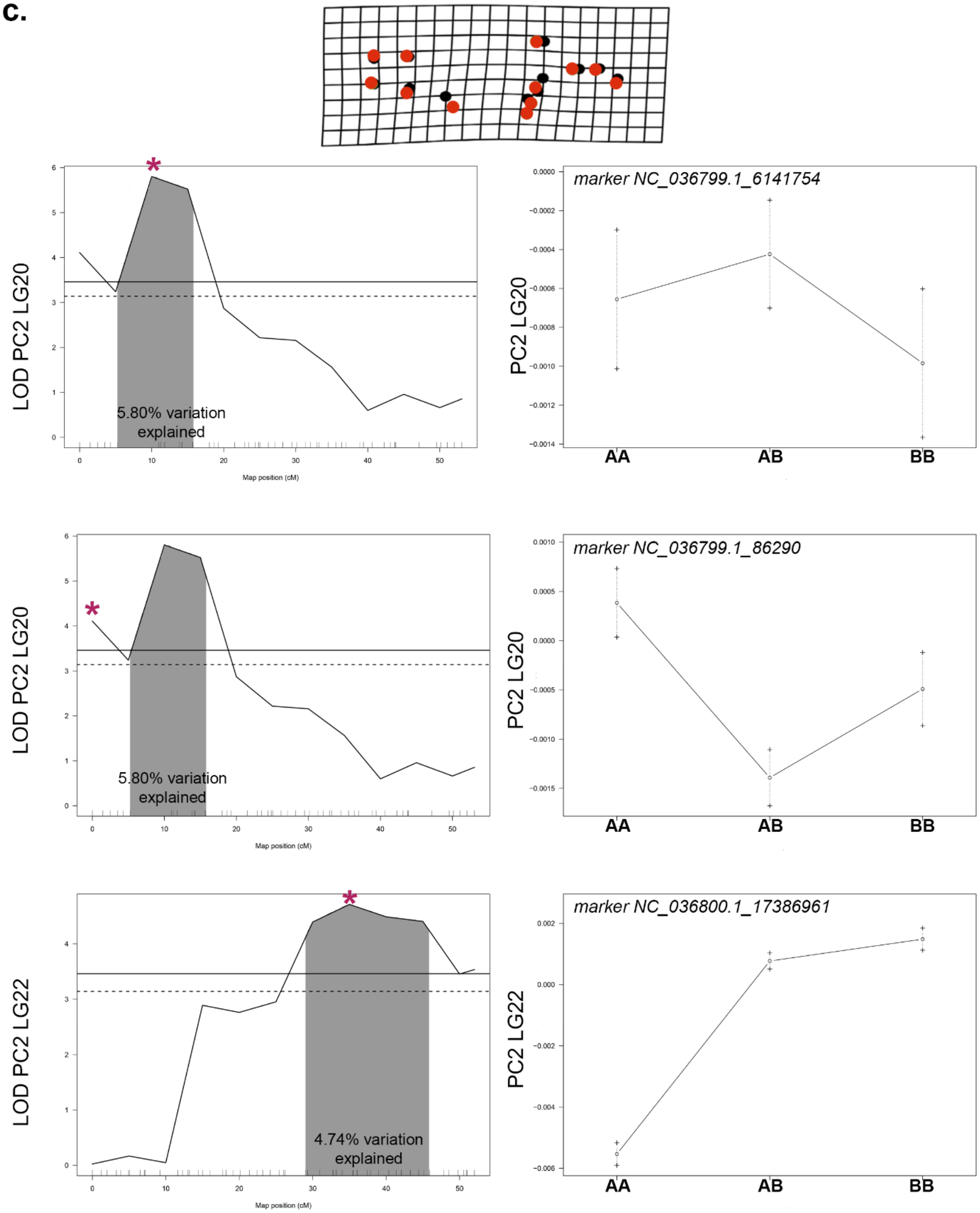

LOD PC2 LG20
5.80% variation explained
Map position (cM)
marker NC_036799.1_6141754
PC2 LG20
AA
AB
BB
LOD PC2 LG20
5.80% variation explained
Map position (cM)
marker NC_036799.1_86290
PC2 LG20
AA
AB
BB
LOD PC2 LG22
4.74% variation explained
Map position (cM)
marker NC_036800.1_17386961
PC2 LG22
AA
AB
BB

**d.**

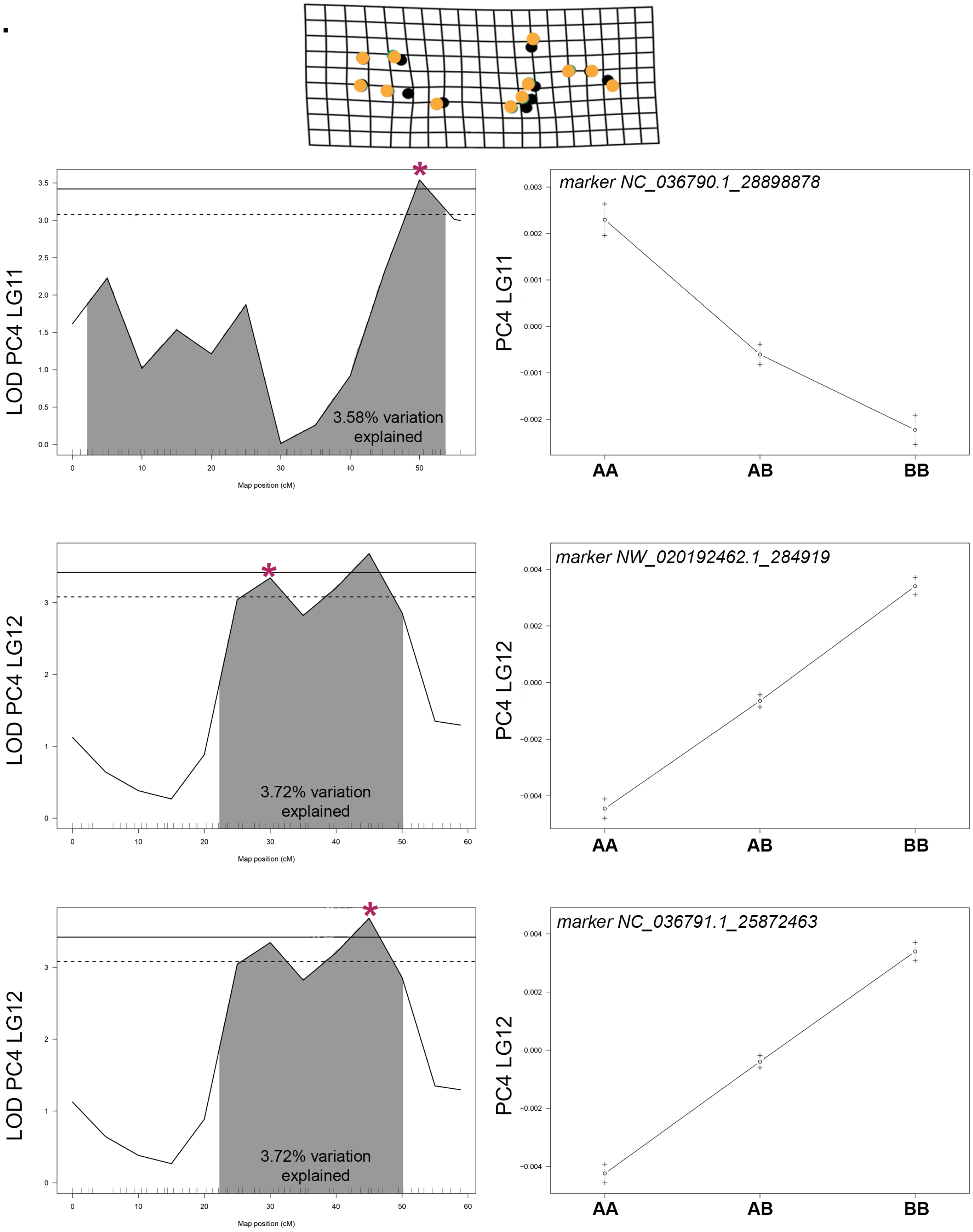

LOD PC4 LG11
3.58% variation explained
Map position (cM)
marker NC_036790.1_28898878
PC4 LG11
AA
AB
BB
LOD PC4 LG12
3.72% variation explained
Map position (cM)
marker NW_020192462.1_284919
PC4 LG12
AA
AB
BB
LOD PC4 LG12
3.72% variation explained
Map position (cM)
marker NC_036791.1_25872463
PC4 LG12
AA
AB
BB

**e.**

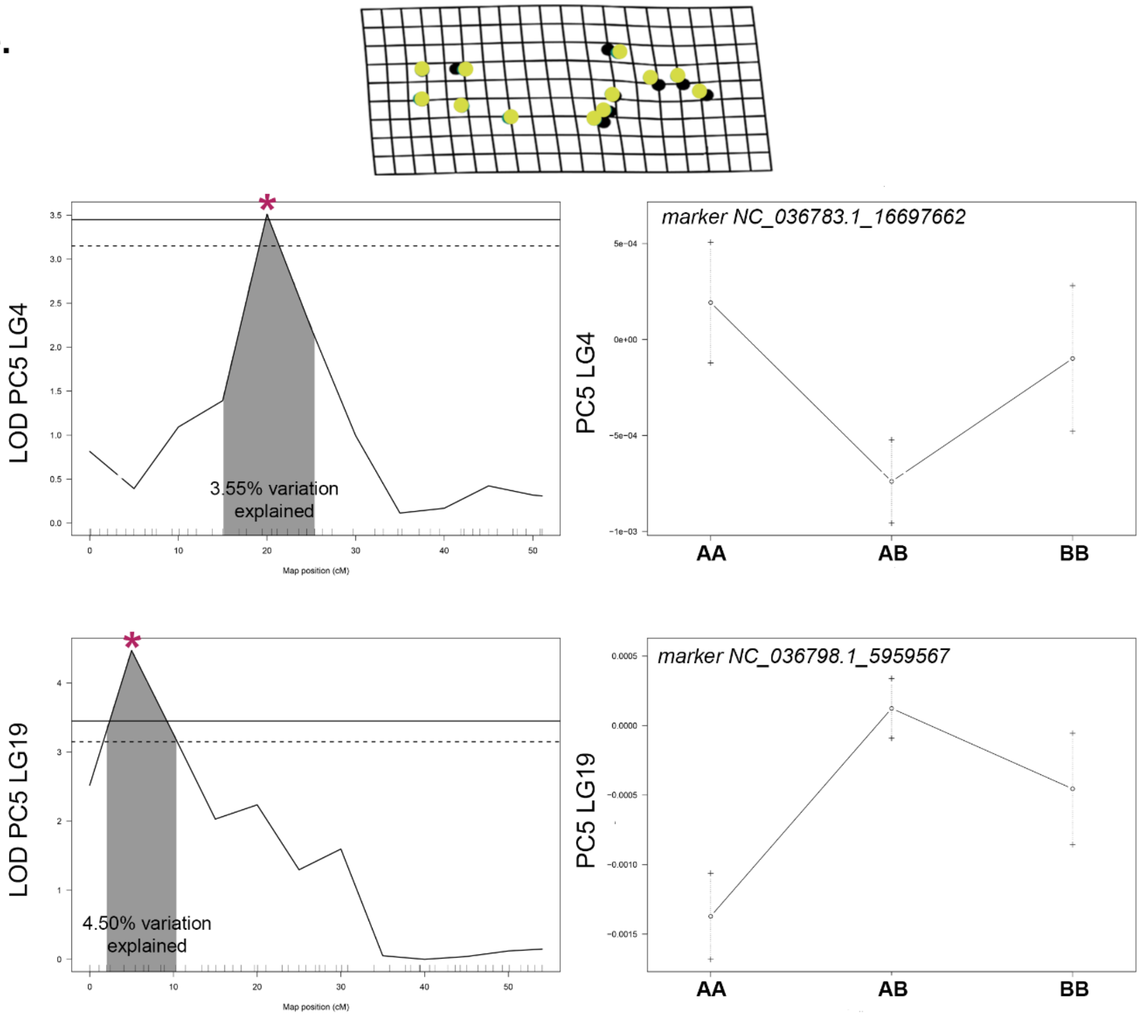

LOD PC5 LG4
3.55% variation explained
Map position (cM)
marker NC_036783.1_16697662
PC5 LG4
AA
AB
BB
LOD PC5 LG19
4.50% variation explained
Map position (cM)
marker NC_036798.1_5959567
PC5 LG19
AA
AB
BB

**f.**

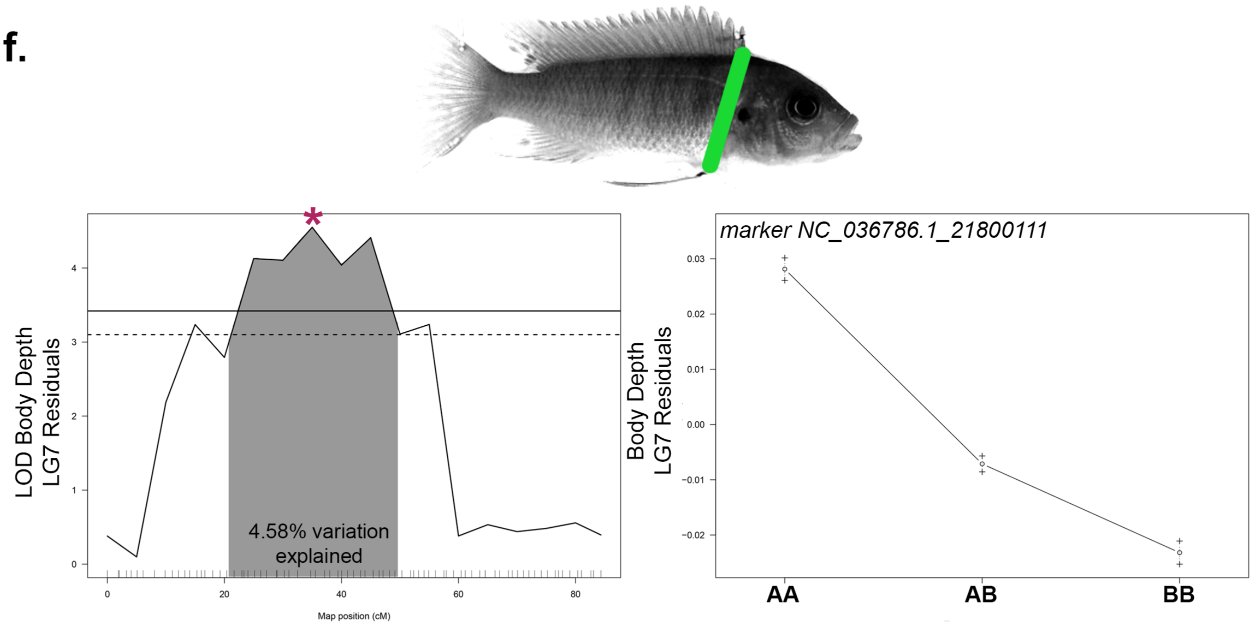

LOD Body Depth
LG7 Residuals
4.58% variation
explained
Map position (cM)
marker NC_036786.1_21800111
Body Depth
LG7 Residuals
AA
AB
BB

g.

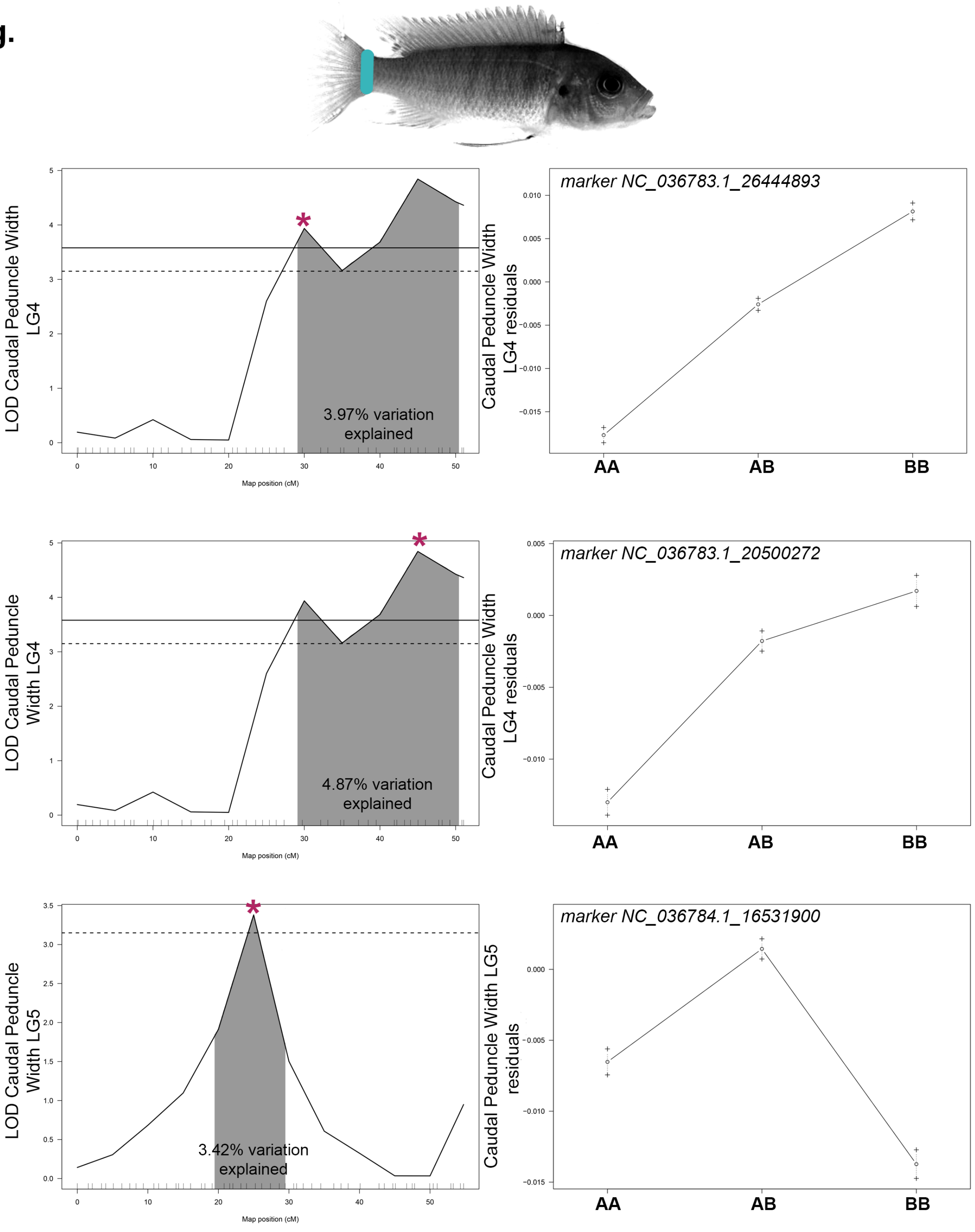
LOD Caudal Peduncle Width LG4
3.97% variation explained
Map position (cM)
marker NC_036783.1_26444893
Caudal Peduncle Width LG4 residuals
AA
AB
BB
LOD Caudal Peduncle Width LG4
4.87% variation explained
Map position (cM)
marker NC_036783.1_20500272
Caudal Peduncle Width LG4 residuals
AA
AB
BB
LOD Caudal Peduncle Width LG5
3.42% variation explained
Map position (cM)
marker NC_036784.1_16531900
Caudal Peduncle Width LG5 residuals
AA
AB
BB

g.

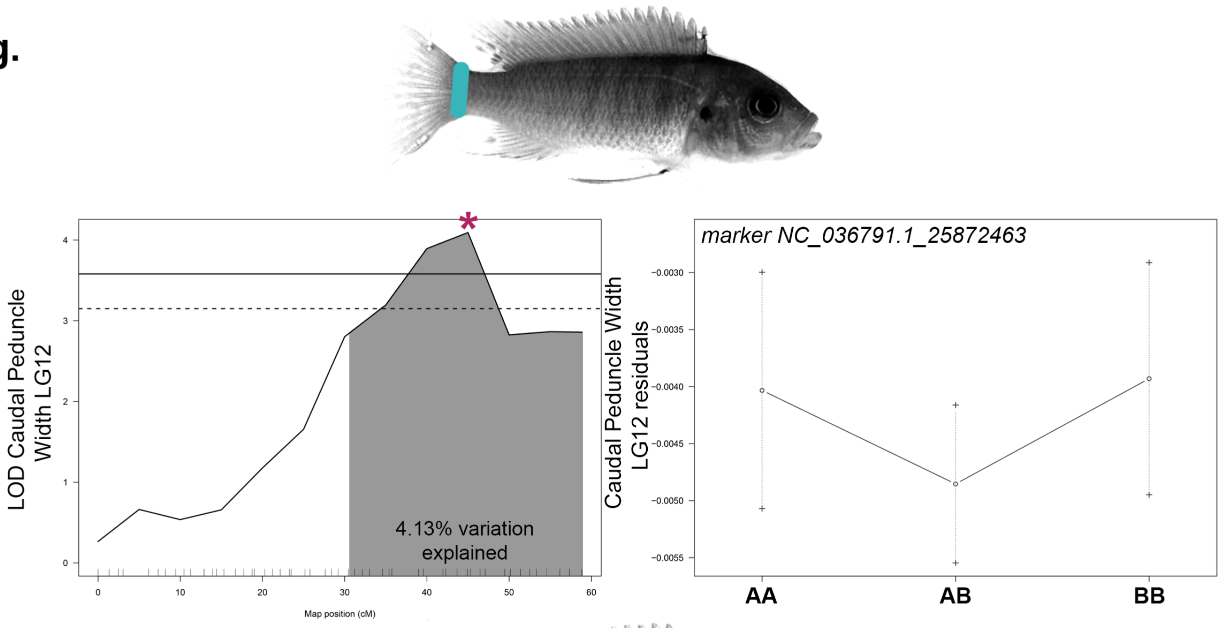
LOD Caudal Peduncle Width LG12
4.13% variation explained
Map position (cM)
marker NC_036791.1_25872463
Caudal Peduncle Width LG12 residuals
AA
AB
BB

h.

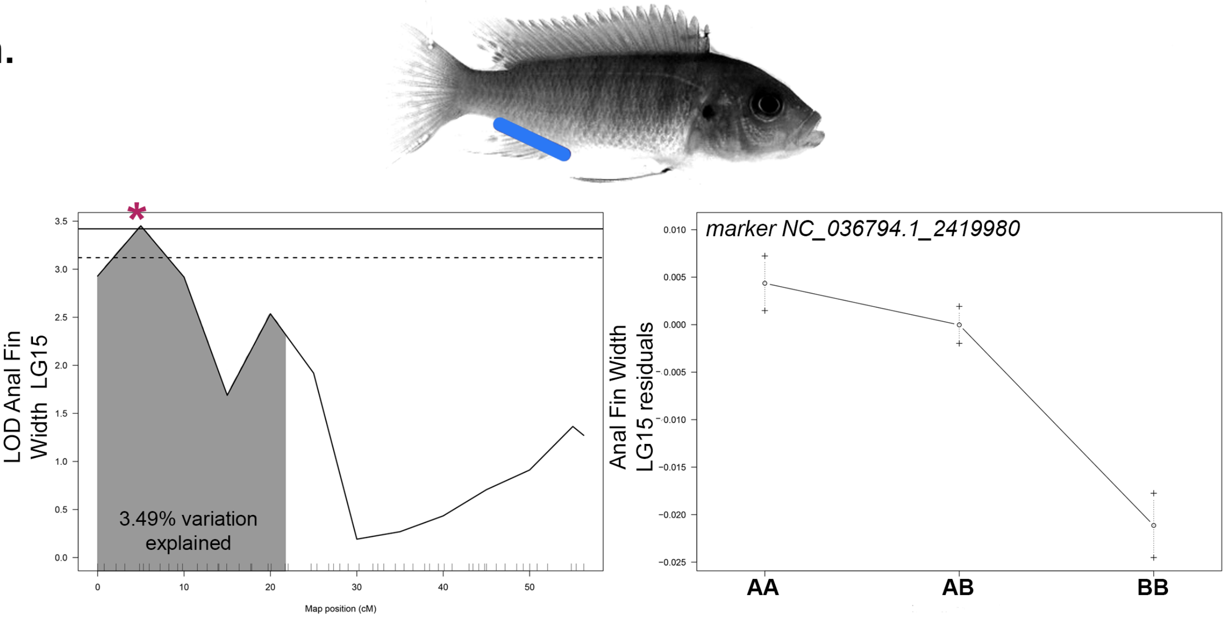
LOD Anal Fin Width LG15
3.49% variation explained
Map position (cM)
marker NC_036794.1_2419980
Anal Fin Width LG15 residuals
AA
AB
BB

i.

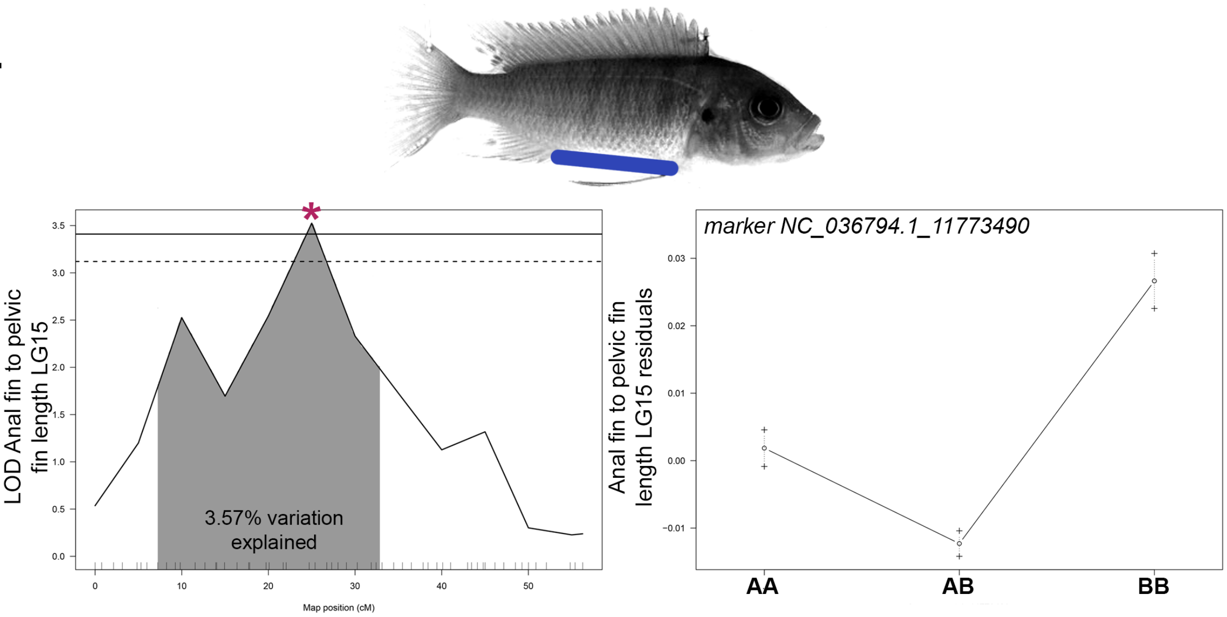


j.

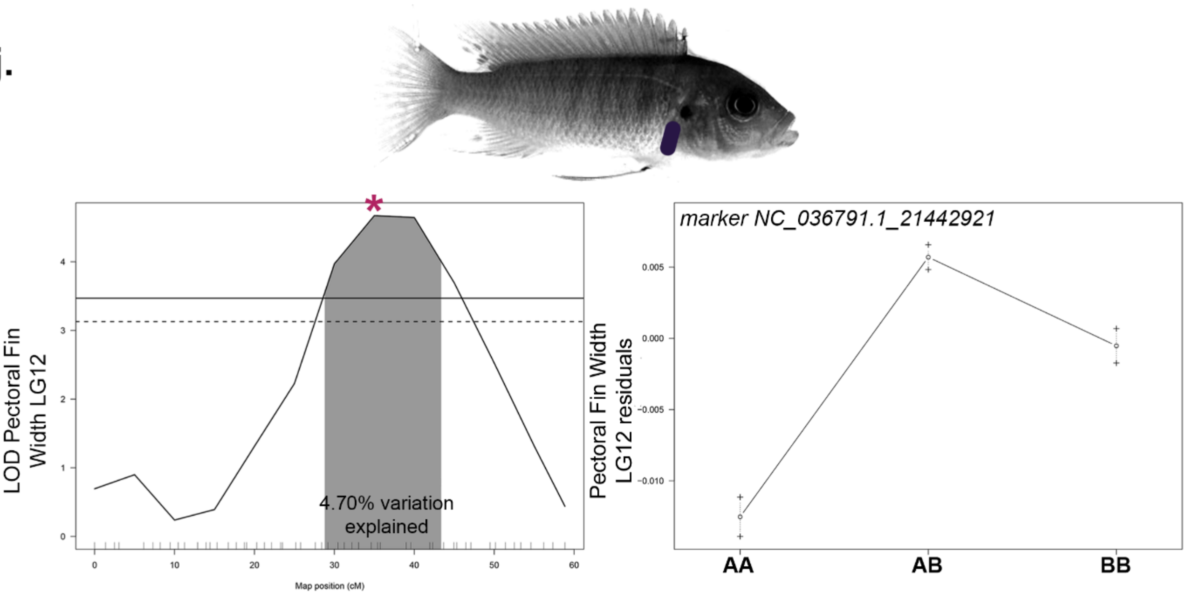


**Figure S5. QTL scans and allelic effects for significant QTL in *Labidochromis* x *Labeotropheus* cross.** Phenotypic measures are indicated by illustration with colors matching Figures 2 and S3 and include (a) standard length, (b) PC1 shape, (c) PC2 shape, (d) PC4 shape, (e) PC5 shape, (f) body depth, (g) caudal peduncle width, (h) anal fin width, (i) length between the anal and pelvic fins, and (j) pectoral fin width. PC3, head proportion, and length between the caudal and anal fins are not included in this figure as there were not any significant or suggestive QTL for these traits in this cross. 95% confidence interval for QTL is indicated by shading, percent of total phenotypic variation explained by QTL is reported, and genome-wide significance is shown at the 5% (solid line) and 10% (dashed line) level. Details of QTL scan are in Table S1 and genome-wide visuals are in Figure S2. Allelic effects are shown for the position indicated by *. The A allele was inherited from the *Labidochromis* granddam and the B allele from the *Labeotropheus* grandsire.
